# Novel genes required for surface-associated motility in ***Acinetobacter baumannii***

**DOI:** 10.1101/2020.03.18.992537

**Authors:** Ulrike Blaschke, Evelyn Skiebe, Gottfried Wilharm

## Abstract

*Acinetobacter baumannii* is an opportunistic and increasingly multi-drug resistant human pathogen rated as a critical priority 1 pathogen for the development of new antibiotics by the WHO in 2017. Despite the lack of flagella, *A. baumannii* can move along wet surfaces in 2 different ways: via twitching motility and surface-associated motility. While twitching motility is known to depend on type IV pili, the mechanism of surface-associated motility is poorly understood. In this study we established a library of 30 *A. baumannii* ATCC 17978 mutants that displayed deficiency in surface-associated motility. By making use of natural competence we also introduced these mutations into strain 29D2 to differentiate strain-specific versus species-specific effects of mutations. Mutated genes were associated with purine/pyrimidine/folate biosynthesis (e.g. *purH*, *purF*, *purM*, *purE*), alarmone/stress metabolism (e.g. Ap4A hydrolase), RNA modification/regulation (e.g. methionyl-tRNA synthetase), outer membrane proteins (e.g. *ompA*), and genes involved in natural competence (*comEC*). All tested mutants originally identified as motility-deficient in strain ATCC 17978 also displayed a motility-deficient phenotype in 29D2. By contrast, further comparative characterization of the mutant sets of both strains regarding pellicle biofilm formation, antibiotic resistance, and virulence in the *Galleria mellonella* infection model revealed numerous strain-specific mutant phenotypes. Our studies highlight the need for comparative analyses to characterize gene functions in *A. baumannii* and for further studies on the mechanisms underlying surface-associated motility.

## Introduction

*Acinetobacter baumannii* is a Gram-negative and strictly aerobic coccobacillus [1, 2]. Being an opportunistic human pathogen [3], *A. baumannii* is associated with nosocomial diseases including soft tissue, bloodstream, and urinary tract infections as well as pneumonia [2]. Worldwide, about 9% of culture-positive infections found in intensive care units arise from *Acinetobacter spp.* [4]. Increased multi-drug resistance in *A. baumannii* has become problematic in recent years [5, 6]. A global surveillance study found that 44% of 18,741 collected isolates were multi-drug resistant. During the study period the proportion of multi-drug resistant *A. baumannii* isolates increased from 23% in 2004 to 63% in 2014 [7]. As a consequence of rising multi-drug resistance, *A. baumannii* was rated as one of the critical priority 1 pathogens for the development of new antibiotics by the WHO in 2017 [8]. Drug resistance and environmental persistence have enabled *A. baumannii* to successfully establish in the hospital environment. Some clinical isolates can survive 100 days or more under dry conditions [9–13]. An important factor for the interaction of *A. baumannii* with biotic or abiotic surfaces is the formation of biofilms, a feature that is associated with an increased tolerance to desiccation stress [14].

A connection between *A. baumannii* virulence and motility has been shown in the *Caenorhabditis elegans* infection model where hypermotility resulted in increased virulence [15]. Although *A. baumannii* does not produce flagella, it is capable of moving in two different ways: via twitching motility and surface-associated motility. For *A. baumannii*, twitching motility has been shown to depend on type IV pili (T4P) [16, 17] which drive the bacteria via retraction of attached T4P [18–25]. Inactivation of the putative T4P retraction ATPase *pilT* reduces twitching motility [26-28,11] but does not abolish surface-associated motility [26, 16]. Surface-associated motility in *A. baumannii* occurs at the surface of semi-dry media and is independent of T4P [26, 29]. Surface-associated motility is poorly understood mechanistically, but was demonstrated to be controlled by quorum sensing [26], light [30], and iron availability [31, 32]. Also, the synthesis of 1,3-diaminopropane (DAP) [33] and lipopolysaccharide (LPS) production [32] were shown to contribute to surface-associated motility of *A. baumannii*. Several genes have been identified which contribute to *A. baumannii*’s capacity for surface-associated motility [26, 32], including a ribonuclease T2 family protein [34] and the superoxide dismutase SodB [35]. A recent study revealed the regulatory control of surface-associated motility and biofilm formation by a cyclic-di-GMP signaling network in *A. baumannii* strain ATCC 17978 [36]. Interestingly, studies on phase-variable phenotypes in *A. baumannii* strain AB5075 showed that “opaque phase” bacterial colonies had improved surface-associated motility [37, 38]. A correlation between pellicle biofilm formation and surface-associated motility has been described in *A. baumannii* [39]. Given the fact that many *A. baumannii* clinical isolates exhibit surface-associated motility, it could be an important trait associated with infection [33,28,26].

To investigate the mechanisms underlying surface-associated motility, we utilized a previously generated transposon mutant library of ATCC 17978 [33] which we screened for a surface-associated motility-deficient phenotype. The motility-deficient mutations were found to affect purine/pyrimidine/folate biosynthesis, alarmone/stress metabolism, RNA modification/regulation, outer membrane proteins, and DNA modification. We characterized these mutants with respect to growth, pellicle biofilm formation, antibiotic resistance, and virulence in the *Galleria mellonella* infection model. To facilitate distinguishing between strain-specific and species-specific traits some mutations were also introduced into the naturally competent *A. baumannii* strain 29D2 [40].

## Materials and Methods

### Bacterial strains and culture conditions

*A. baumannii* strain ATCC 17978 was purchased from LGC Promochem. The *A. baumannii* strain 29D2 was isolated from a white stork [40] and is naturally competent [41]. All strains were grown at 37°C in Luria-Bertani (LB) broth or on LB agar, and mutants were supplemented with 50 µg/mL of kanamycin. All strains used in this work are listed in supplementary Table S1. Single colonies were used as inoculum for overnight cultures or motility plates. Neither strain ATCC 17978 nor strain 29D2 exhibited phase variation [38,37,42].

### Bacterial transformation and generation of an *A. baumannii* mutant library

ATCC 17978 transposon mutants were generated using the EZ-Tn5™ <KAN-2> Insertion Kit (Epicentre Biotechnologies) as previously described [33]. Transformation of the transposome complex into ATCC 17978 was performed by electroporation [43]. 29D2 mutants were generated by making use of the strain’s ability for natural competence. The transforming DNA was isolated from the ATCC 17978 mutants described above. A suspension of DNA-accepting bacteria was generated by resuspending a few colonies in 100 µL of sterile PBS. The bacterial suspension was then mixed with equal volumes of the transforming DNA (∼400 ng/µL). This mixture was stabbed into motility agar plates 10 times, pipetting 2 µL of the mixture with each stabbing [16]. The motility plates were incubated for 18 h at 37°C. After incubation, the bacteria were flushed off the motility plates with 1 mL of sterile PBS and 100 µL was plated on selective agar plates (50 g/mL of kanamycin). After sub-culturing of selected colonies transformation was confirmed by PCR.

### Identification of transposon insertion sites by single-primer PCR

To identify the transposon insertion sites of ATCC 17978 motility mutants, single-primer PCR was performed as described previously [33] using one of the following primers: FP-2Kana 5’-CTTCCCGACAACGCAGACCG-3’; FP-3Kana 5’-GAGTTGAAGGATCAGATCACGC-3’; RP-2Kana 5’-CCCTTGTATTACTGTTTATGTAAGC-3’; RP-3Kana 5’-CGCGGCCTCGAGCAAGACG-3’; Tn5-Kana-For4 5’-GTTTTCTCCTTCATTACAGAAACG-3’; and Tn5-Kana-Rev4 5’-

CCCATACAATCGATAGATTGTCG-3’. Transposon insertions of all mutants (ATCC 17978 and 29D2) were confirmed by PCR using primers for the EZ-Tn5™ <KAN-2> kanamycin cassette (Suppl. Fig. S1), which are specified in the manufacturer’s instructions, and appropriate gene target site primers (Suppl. Table S2; Suppl. Figs. S2 and S3).

### Surface-associated motility

Motility assays were performed as described previously [33]. A single bacterial colony from a nutrient agar plate (Oxoid) or selective agar plates (supplemented with 50 µg/mL of kanamycin for the mutants) of either wildtype (ATCC 17978 and 29D2) or mutants was lifted with a pipette tip and transferred to the surface of a motility plate (0.5% agarose). Plates were incubated for 16 h at 37°C. The diameter of the surface motility spreading zone was measured and quadruplicates were statistically analyzed.

### Bacterial growth curves

Growth curves were determined by growing overnight cultures at 37°C in LB medium (supplemented with 50 µg/mL of kanamycin for the mutants). Overnight cultures were adjusted to 1 OD (600 nm) in LB medium. In 250 mL baffled flasks, 50 mL of LB medium (without antibiotics) was inoculated with 1 mL of the OD-adjusted inoculum. The cultures were incubated at 37°C for 9 h with shaking at 160 rpm. OD measurements at 600 nm were performed every hour by sampling 100 µL of every culture. For each strain, data obtained from 3 independent cultures grown on the same day were averaged and represented by the mean ± SD.

### Infection in the *Galleria mellonella* caterpillar

For *G. mellonella* caterpillar infection, bacteria were grown in LB medium overnight at 37°C (50 µg/mL of kanamycin was added to mutant strains). Cultures were diluted 1:50 in LB medium and incubated for another 4 h at 37°C. Bacteria were pelleted for 5 min at 7500 rpm at room temperature (RT) and the supernatant was discarded. Bacteria were resuspended in 500 µL sterile PBS, adjusted to an OD_600_ nm of 1.0 and diluted 1:10 in sterile PBS. 5 µL of this dilution, corresponding to 3 x 10^5^ colony-forming units (CFUs), was injected into the last right proleg of *G. mellonella* caterpillars (purchased from TZ-TERRARISTIK, Germany, and BioSystems Technology TruLarv, UK). As a control, caterpillars were injected with 5 µL of sterile PBS. Three independent experiments were performed with groups of 16 caterpillars for every bacterial strain and control. The caterpillars were incubated at 37°C for 5 days and checked daily for vitality. Experiments with more than 2 dead caterpillars within 5 days in the control group were not considered valid. CFUs were determined by serial dilutions, plated on nutrient agar, and colonies were counted after incubation at 37°C for 18 h. For each strain, data obtained from 3 independent experiments were averaged and represented by the mean ± SD.

### Determination of susceptibility to antibiotics

For the minimal inhibitory concentration (MIC) tests, bacteria were grown in LB medium overnight at 37°C (to mutant strains 50 µg/mL of kanamycin was added). Cultures were diluted 1:50 in LB medium and incubated (without antibiotics) another 4 h at 37°C. Agar plates were flushed with 2 mL of each culture and E-test strips (Liofilchem, Italy) were deposited on nutrient agar plates. MICs were determined after incubation for 16 h at 37°C. Three independent experiments were performed and statistical significance was tested by the Student’s *t* test (2-tailed, unpaired).

### Pellicle biofilm assays

*A. baumannii* strains were grown in LB medium overnight at 37°C (50 µg/mL of kanamycin was added to mutant strains). The cultures were adjusted to an OD_600_ nm of 1.0 and 3 mL of LB medium (without antibiotics) was inoculated with 15 µL of OD-adjusted culture. Samples were incubated at RT for at least 3 days. The LB medium was removed using a thin cannula and the biofilm (sticking to the tube wall) was stained with a 0.5% crystal violet solution (w/v in Aqua Bidest) for 20 min. The crystal violet was removed and the biofilm was washed twice with 4 mL Aqua Bidest. The biofilm was scrubbed and flushed off the tube walls with a pipet tip and 96% alcohol solution. The absorption at 550 nm was determined. Samples which showed an OD > 1.0 were diluted 1:10 with 96% ethanol for measurement. For each strain 3 independent experiments were performed and statistical significance was analyzed by the Student’s *t* test (2-tailed, unpaired).

### Microscopy

The bacterial strains ATCC 17978, ATCC 17978 *ompA::Km*, 29D2, and 29D2 *ompA::Km* were grown for 16 h at 37°C under constant shaking. One μL of each bacterial overnight culture was pipetted on a glass slide and analyzed under the bright field microscope (200 times magnification).

### Statistical analysis

All experiments were performed at least 3 times. Comparison between groups was performed using GraphPad Prism 7 with Student’s *t* test (2-tailed, unpaired). P-values less than 0.05 were considered to be statistically significant.

## Results

### Surface-associated motility

Approximately 2,000 transposon mutants of ATCC 17978 were screened for surface-associated motility phenotypes and 30 were identified with motility defects. Previous studies were limited to the characterization of mutations in single strains. Here, to provide a comparative study, we introduced at least one mutation of every gene function category into 29D2 to get insight into strain-specific and species-specific traits.

To this end, surface-associated motility was analyzed on 0.5% agarose plates. The diameter (Ø) of the surface motility spreading zone of 3 independent experiments was measured and analyzed (Fig. 1A and Suppl. Table S3). All selected motility-deficient mutants of ATCC 17978 exhibited at least a 7-fold reduction of the spreading zone. Subsequently, DNA isolated from these transposon mutants was used to generate mutants in 29D2. All 29D2 mutants displayed a motility-deficient phenotype compared to the wildtype strain (Fig. 1B). Note that the surface-associated motility spreading zone of the wildtype ATCC 17978 (mean Ø of 78 mm) was more than twice as large as that of the 29D2 wildtype strain (mean Ø of 30 mm). Most ATCC 17978 mutants showed a 16-fold reduced surface-associated motility compared to the wildtype strain (Fig 2A), whereas the *a1s_0806* (encoding an aminotransferase) mutant lacked almost any measurable surface-associated motility (mean Ø of 1 mm). 3 mutants, *purH::Km* (mean Ø 10.25 mm), *1970::Km* (mean Ø 8.75 mm), and *3297::Km* (mean Ø 11 mm), showed 10-fold reduced surface-associated motility. Most 29D2 mutants displayed a 4-fold reduction in their surface-associated motility. The most pronounced reduction in motility appeared in mutants *purH::Km* (mean Ø 5 mm), *purF::Km* (mean Ø 3.75 mm), and *ddc::Km* (mean Ø 4.25 mm). The mutant *purM::Km* (mean Ø of about 16 mm) had the lowest reduction in surface-associated motility.

**Fig. 1.**
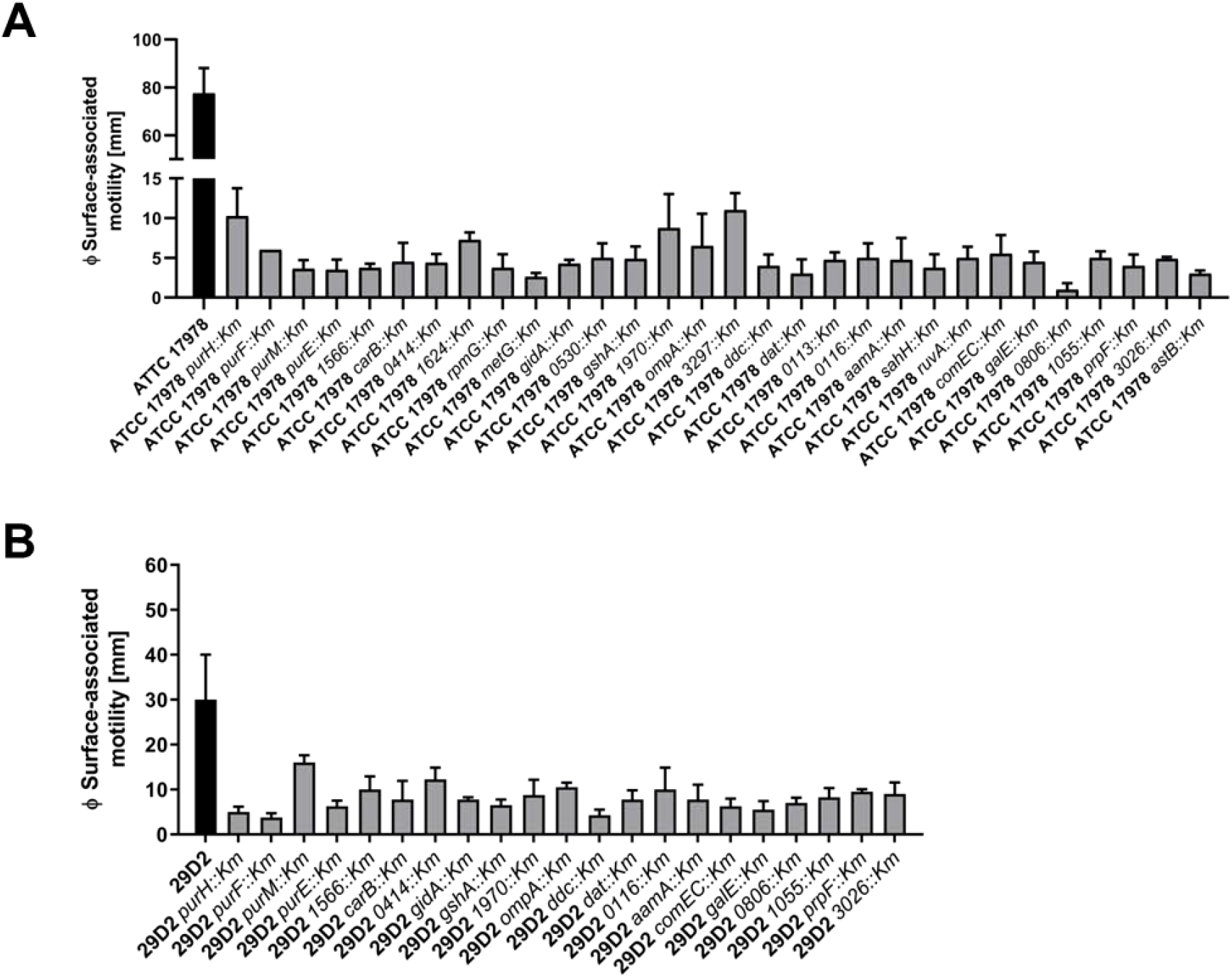
ATCC 17978 mutants (A) and 29D2 mutants (B) deficient in surface-associated motility. Wildtypes and mutants of strains ATCC 17978 and 29D2 were inoculated on motility plates. Plates were incubated for 16 h at 37°C. The diameter (Ø) of the surface-associated motility spreading zone was measured and triplicates were statistically analyzed. All mutants of strains ATCC 17978 (**A**) and 29D2 (**B**) displayed a significant motility deficiency compared to their respective parental strain (p-value ≤ 0.05).

**Fig. 2.**
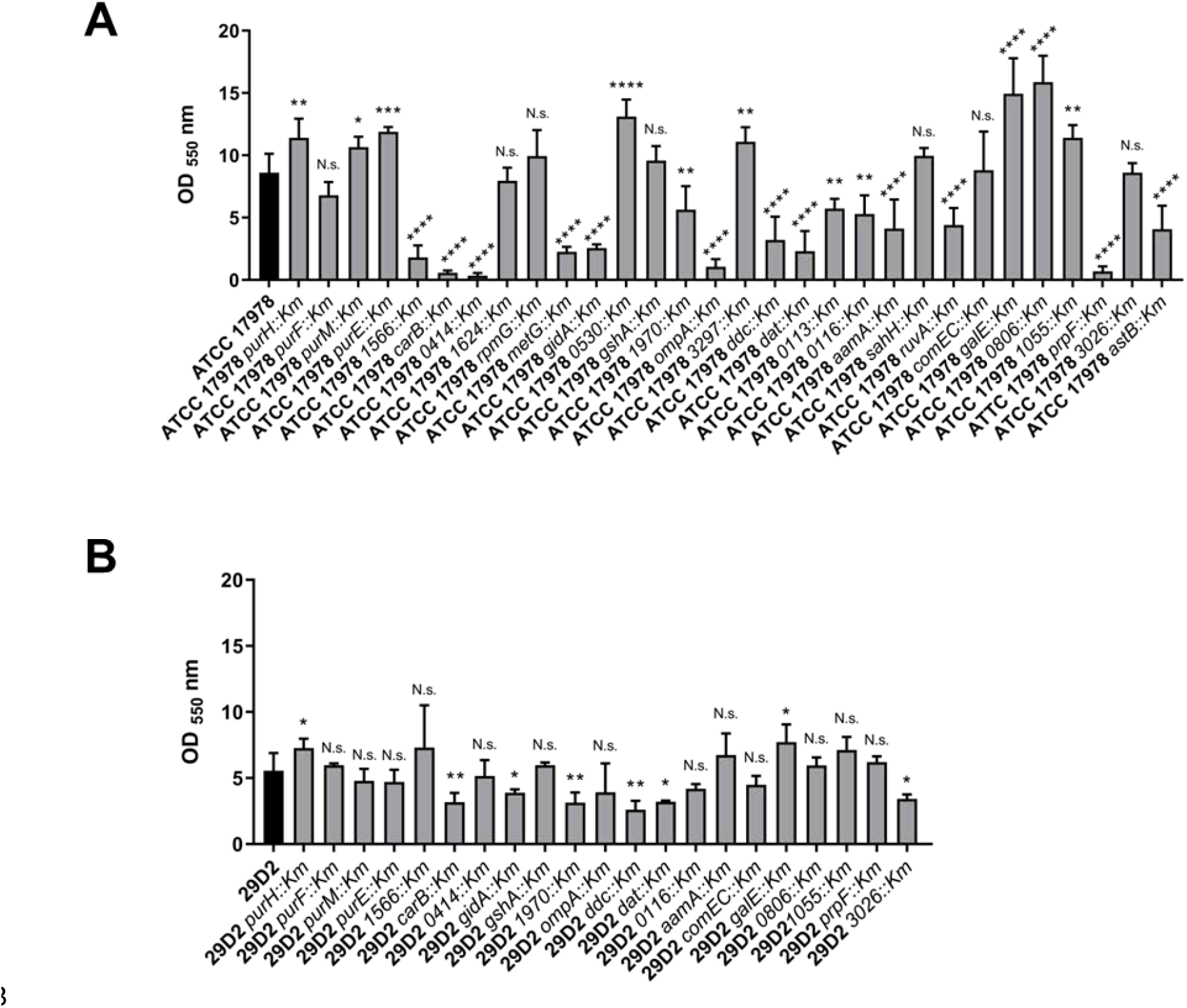
Pellicle biofilm formation of ATCC 17978 wildtype and mutants (A) and 29D2 wildtype and mutants (B). *A. baumannii* pellicle biofilms developed within 3 days of incubation and were stained with a 0.5% crystal violet solution. The biofilm was scrubbed and flushed off the tube walls and the absorption was determined at 550 nm. For each strain 3 independent experiments were performed and statistical significance was analyzed by the Student’s *t* test (2-tailed, unpaired). Significance as indicated: *, p-value ≤ 0.05; **, p-value ≤ 0.01; ***, p-value ≤ 0.001; ****, p-value ≤ 0.0001.

To summarize, all mutations initially identified in ATCC 17978 that conferred motility defects were also found to cause motility-deficient phenotypes when introduced into the orthologous genes of 29D2.

### Pellicle biofilm formation

The formation of pellicles, a specific form of biofilm, occurs at the air-liquid interface and is distinct from submerged biofilms [39,44,45]. A correlation between surface-associated motility and pellicle biofilm formation has been described for *A. baumannii* [39]. We examined the ability of our motility-deficient mutants to form pellicles. Pellicle biofilms were incubated 3 days, stained with a 0.5% crystal violet solution, and analyzed by OD measurements (Suppl. Table S4). Pellicle-biofilm formation in wildtype ATCC 17978 was measured to be about 8.6 at OD_550_ nm (Fig. 2A). A broad spread between low and high pellicle-producing mutants was visible, ranging between a 1.8-fold increase to more than a 25-fold decrease. For 15 of 30 mutants less than 67% of the wildtype-specific pellicle biomass was quantified (Table 1 and Fig. 2A). In the mutants *carB::Km*, *0414::Km,* and *prpF::Km* a pellicle biomass less than 8% compared to the wildtype biomass was measured. This dramatic decrease was not observed by inactivation of the orthologous gene in the 29D2 background. In ATCC 17978, 8 mutants (*purH::Km*, *purM::Km*, *purE::Km*, *0530::Km*, *3297::Km*, *galE::Km*, *0806::Km*, and *1055::Km*) were able to produce more pellicle biomass compared to the wildtype strain, of which the mutants *0530::Km*, *galE::Km,* and *0806::Km* produced 50-80% more pellicle biomass compared to wildtype (Fig. 2A). 29D2 mutants only displayed small changes in pellicle biofilm formation compared to wildtype, with a range of the mutants’ pellicle biomass production from a 1.3-fold increase to a 2.1-fold decrease. 13 of 21 tested 29D2 mutants did not display any significant change in their pellicle biofilm formation compared to the parental strain (Table 2). Deficiencies could be observed in the following 6 mutant strains: *carB::Km*, *1970::Km*, *ddc::Km*, *dat::Km*, *gidA::Km,* and *3026::Km*, which produced less than 70% of the 29D2 wildtype-specific pellicle biomass. Only the following 2 mutants produced significantly more pellicle biomass (Fig. 2B) compared to the wildtype strain: *purH::Km* (30%) and *galE::Km* (38%).

**Table 1.**
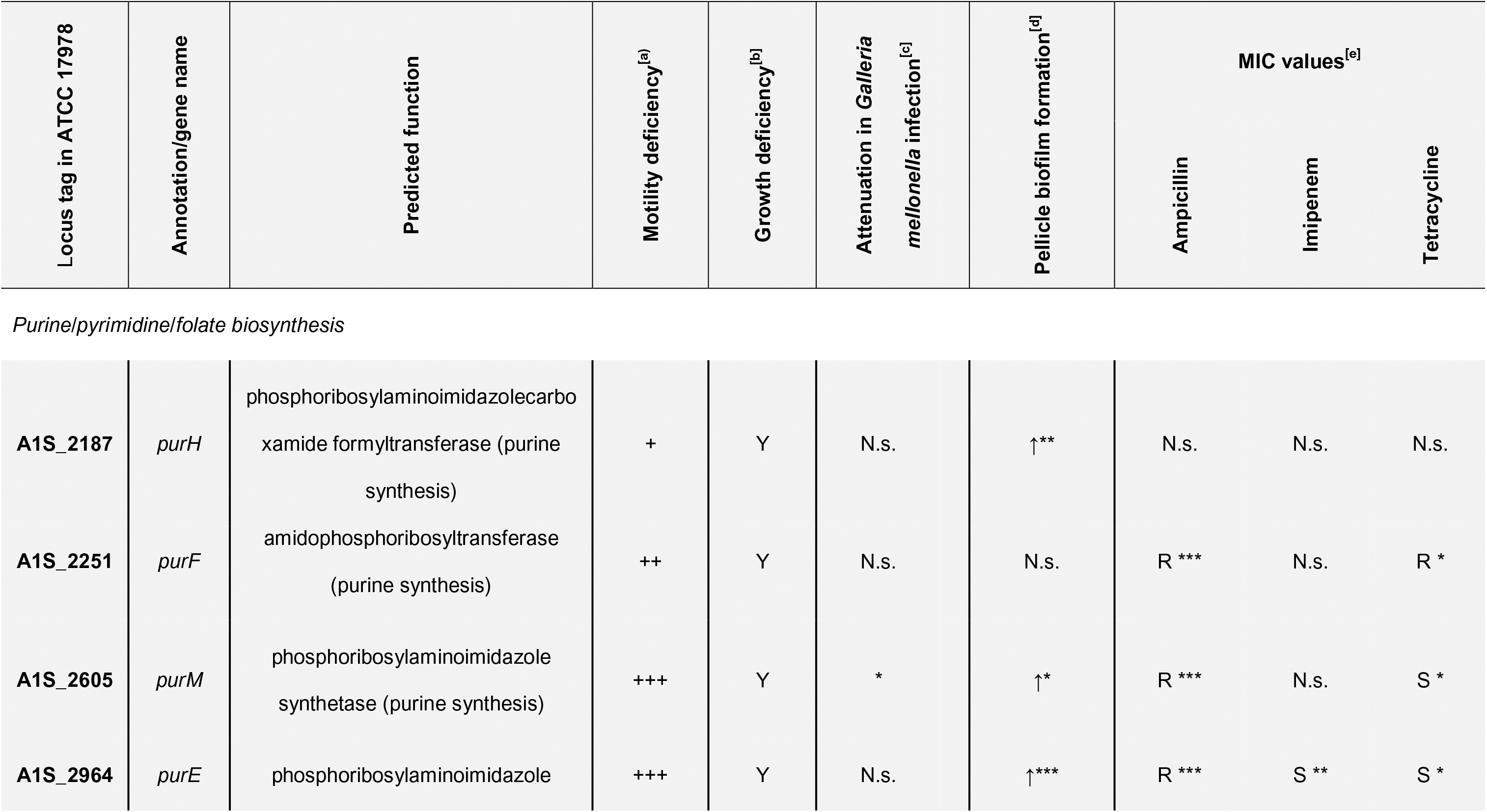
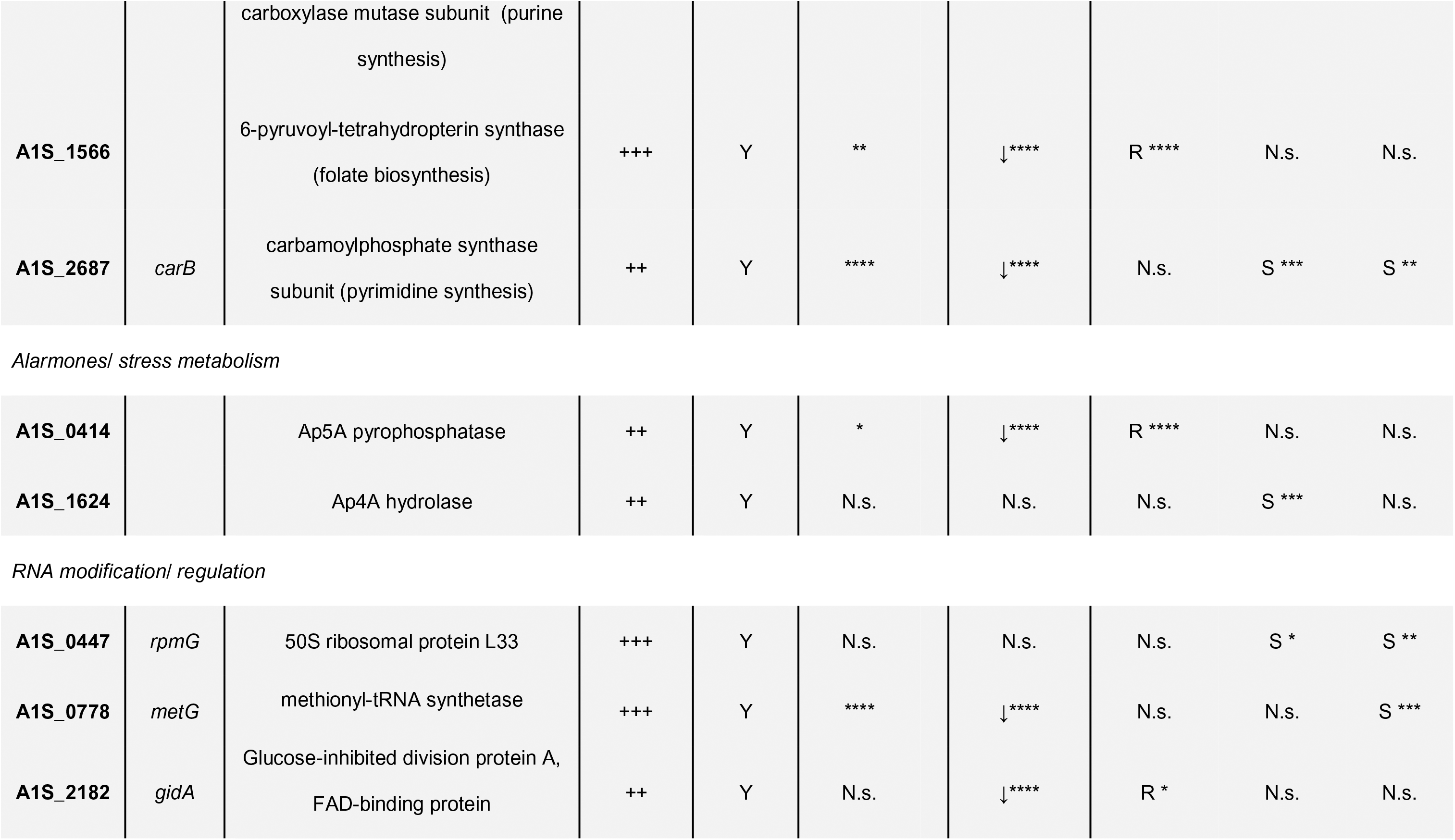
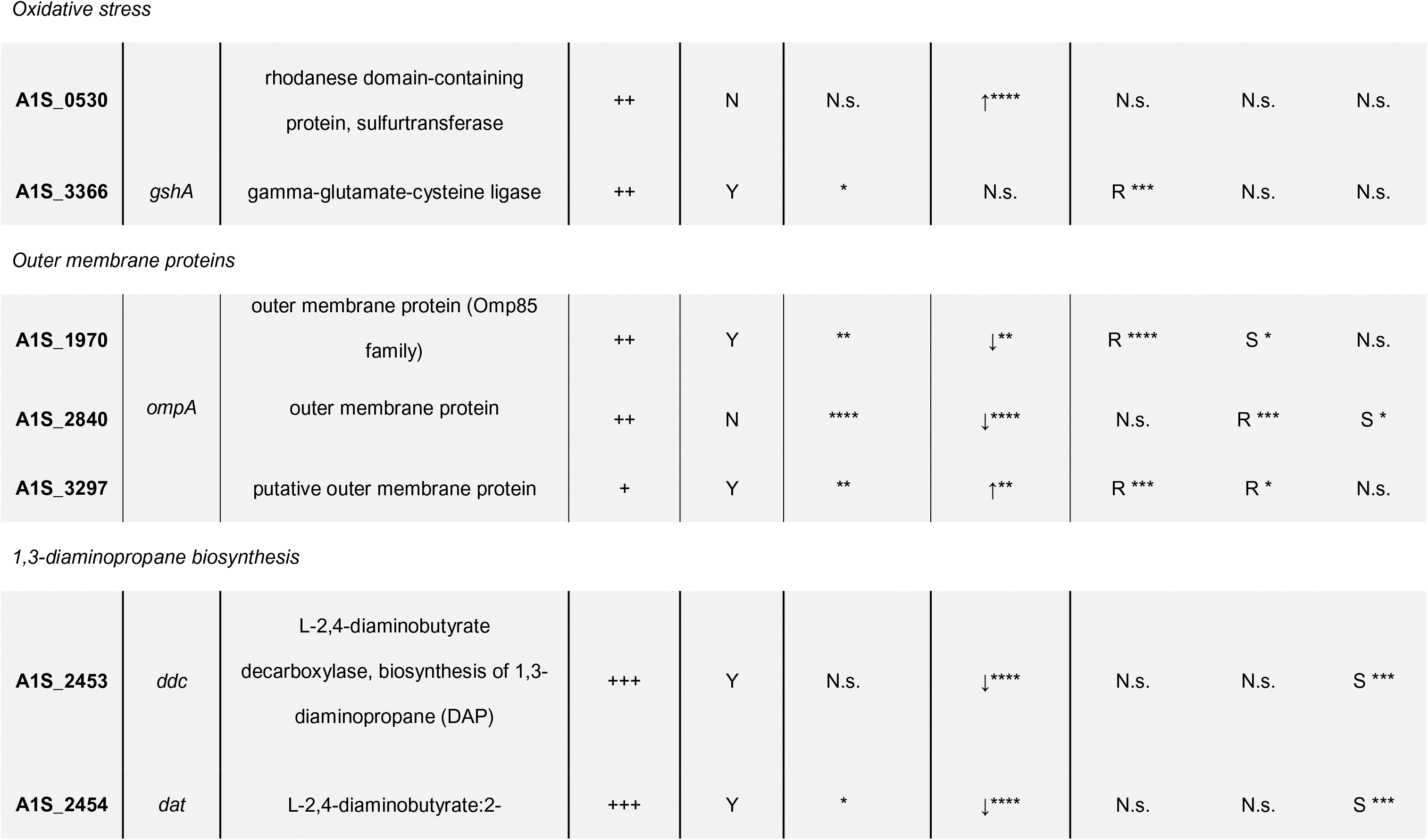
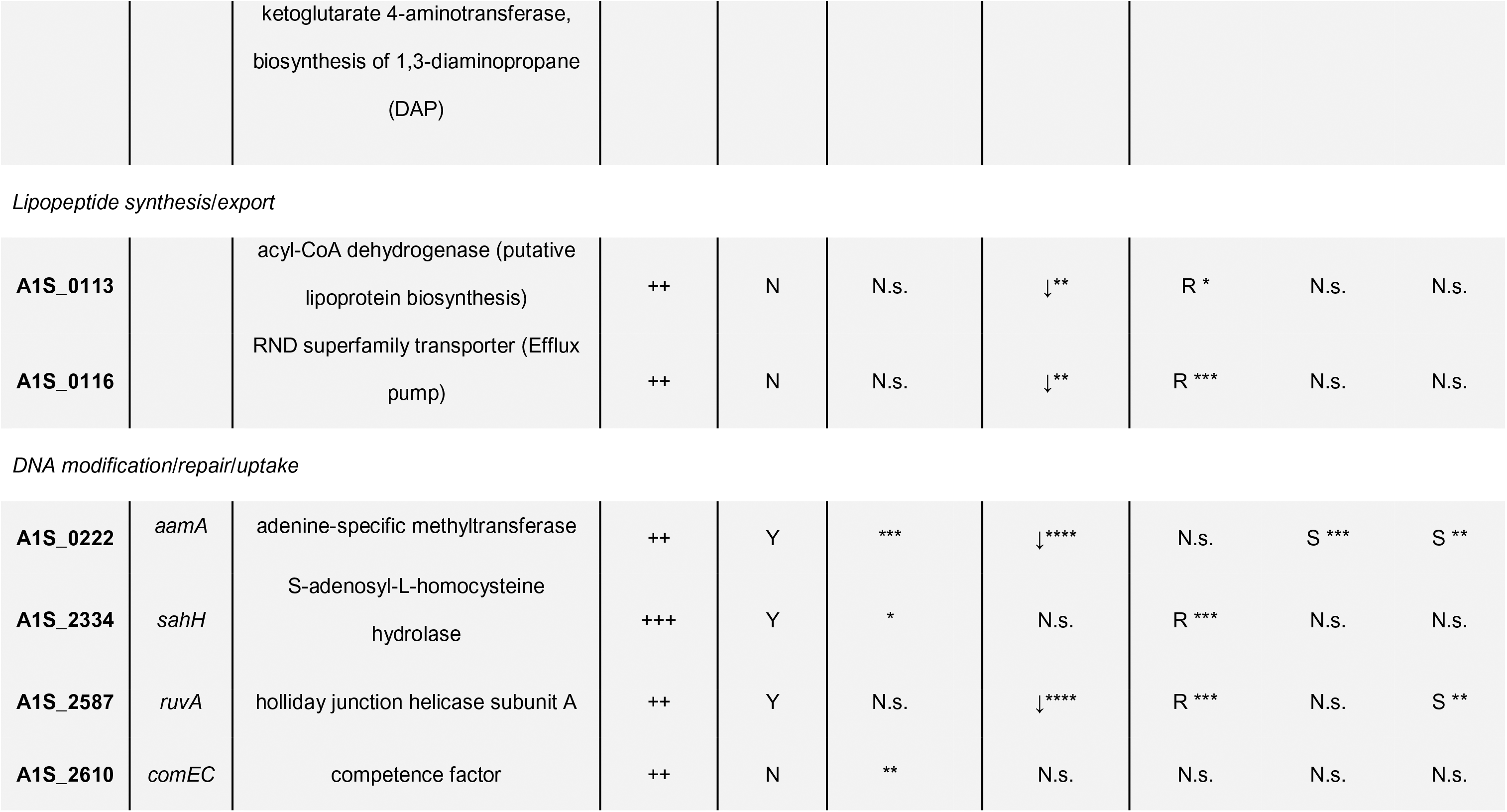
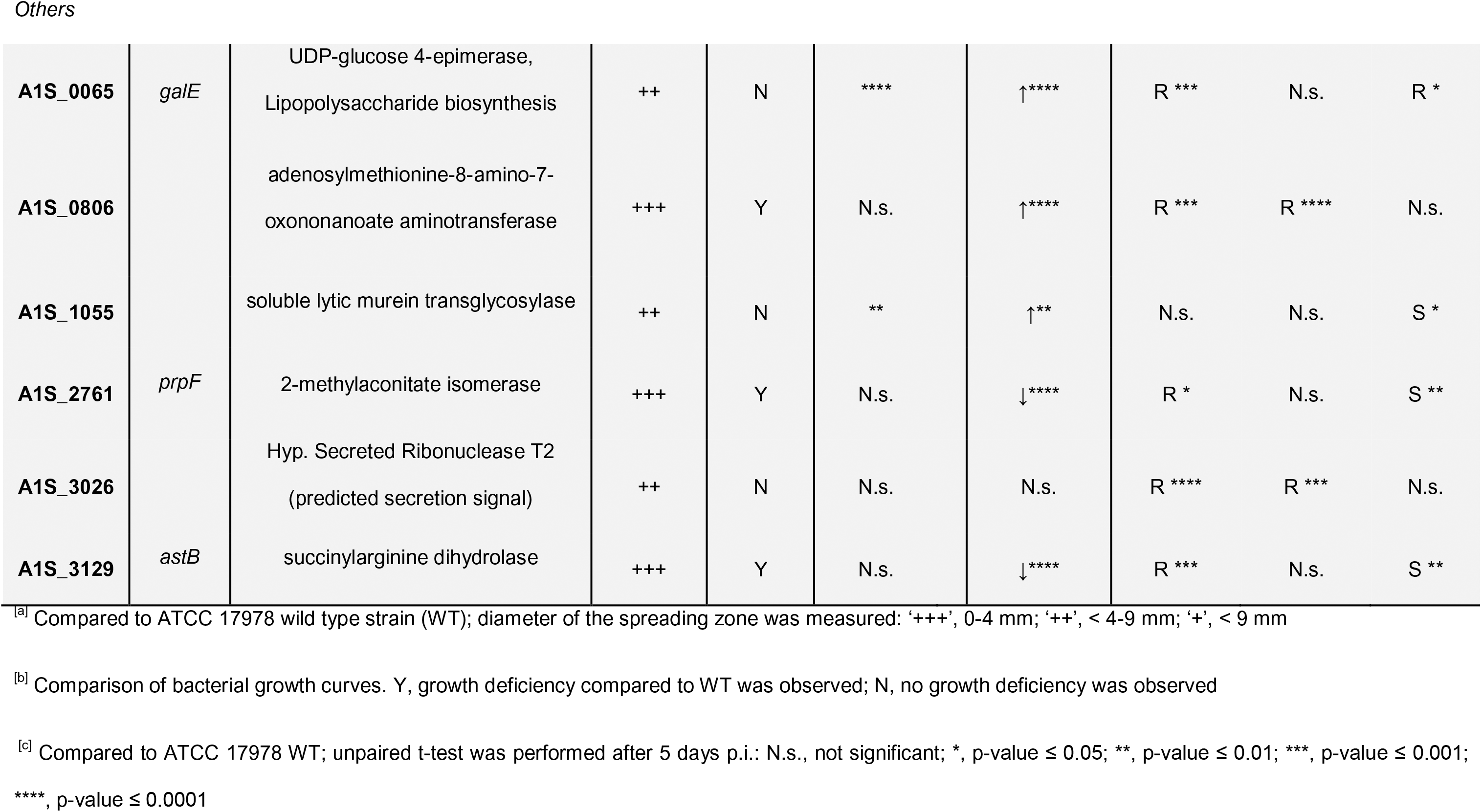
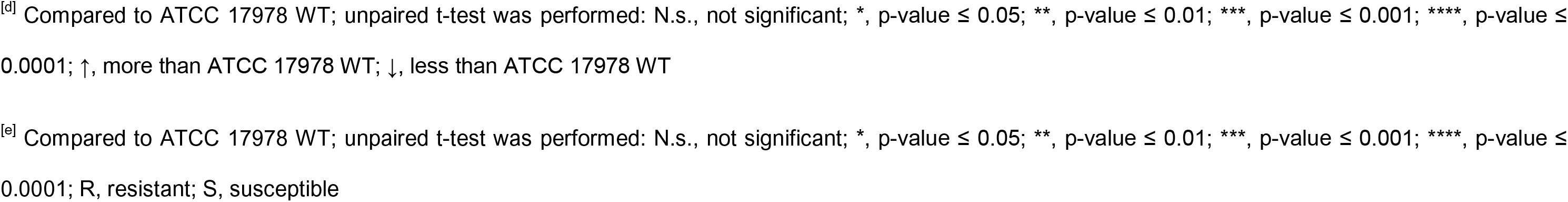
Summary of experimental results on genes involved in surface-associated motility in *A. baumannii* ATCC 17978.

In summary, the ATCC 17978 parental strain produced more pellicle biofilms compared to 29D2. Conspicuous changes in biofilm formation could mainly be observed among ATCC 17978 mutants. Concordance of pellicle formation phenotypes between the mutants of both strains was limited suggesting that strain-specific traits that are independent of surface-associated motility influence pellicle biomass production.

### Bacterial growth

The ability of motility-deficient mutants to grow as a planktonic culture under aeration was assayed. Growth curves and data for all tested strains are provided in supplementary Fig. S4 (ATCC 17978 mutants), supplementary Fig. S5 (29D2 mutants), and Table S5. For 17978, 22 of 30 tested mutant strains exhibited significant growth defects compared to the parental strain (Table 1). The most striking growth defects (Fig. 3A) were observed in the mutants defective in purine biosynthesis (*purH::Km*, *purF::Km*, *purM::Km*, and *purE::Km*), pyrimidine biosynthesis (*carB::Km*), and diaminopropane biosynthesis (*ddc::Km* and *dat::Km*). Only 8 of 30 tested mutant strains were able to grow without any defect compared to the parental strain (Table 1). By testing the 29D2 mutant strains we observed 13 of 21 strains with notable planktonic growth defects (Table 2). Within this group most striking defects were observed with mutations associated with purine biosynthesis (*purH::Km*, *purF::Km*, *purM::Km*, and *purE::Km*), pyrimidine biosynthesis (*carB::Km*), folate biosynthesis (*1566::Km*), and diaminopropane biosynthesis (*ddc::Km* and *dat::Km*). Additionally, *galE::Km*, *comEC::Km,* and *prpF::Km* mutants displayed strong growth deficiencies (Fig. 3B). The mutant *ompA::Km* showed growth comparable to the parental strain for up to 4 h, reached a growth maximum of 2.5 ± 0.28 OD_600_ nm after 5 h, but then slowly collapsed to 1.36 ± 0.73 after 9 h. No growth defects were observed in 8 of 21 tested mutants (Table 2).

**Table 2.**
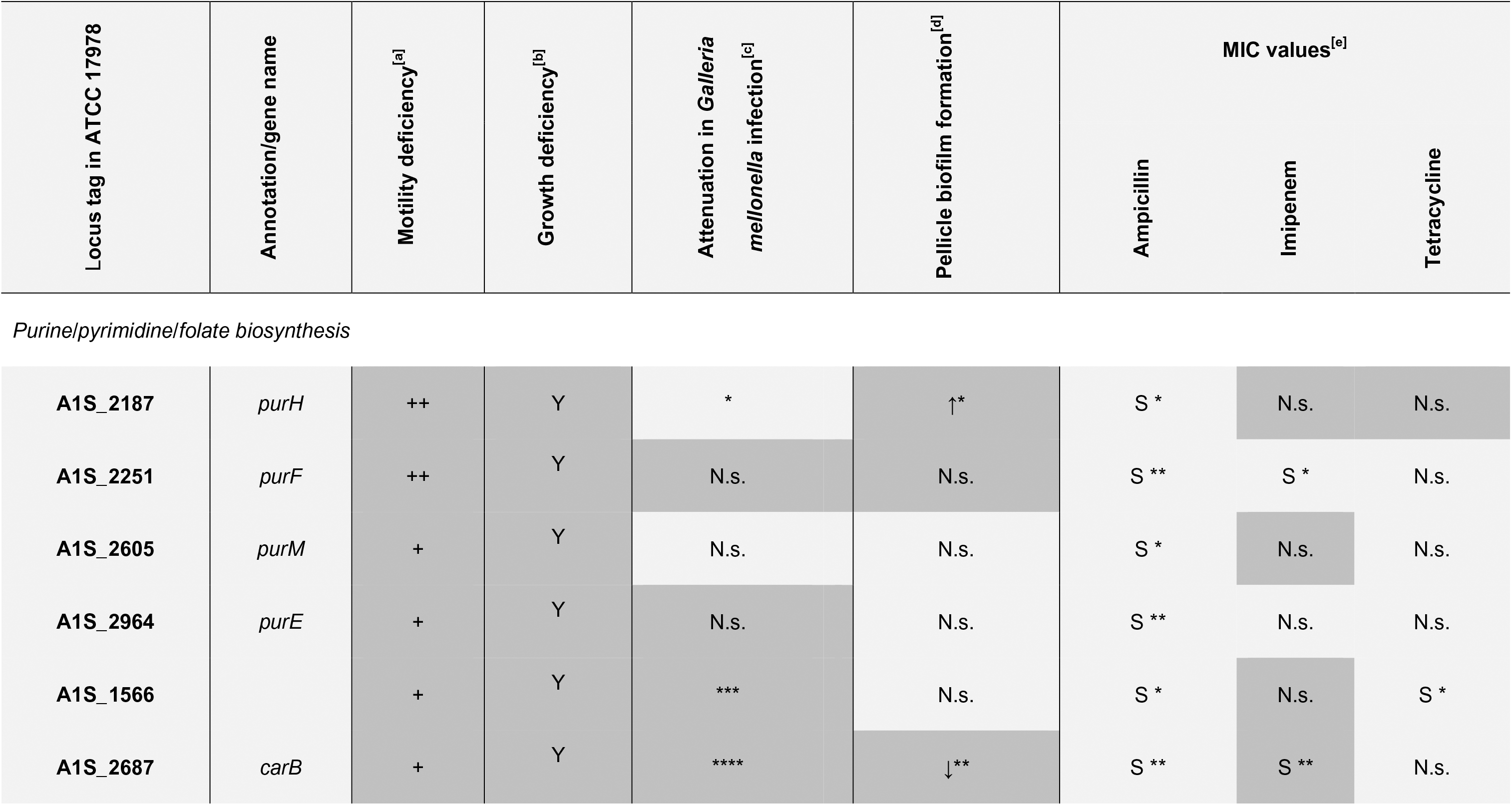
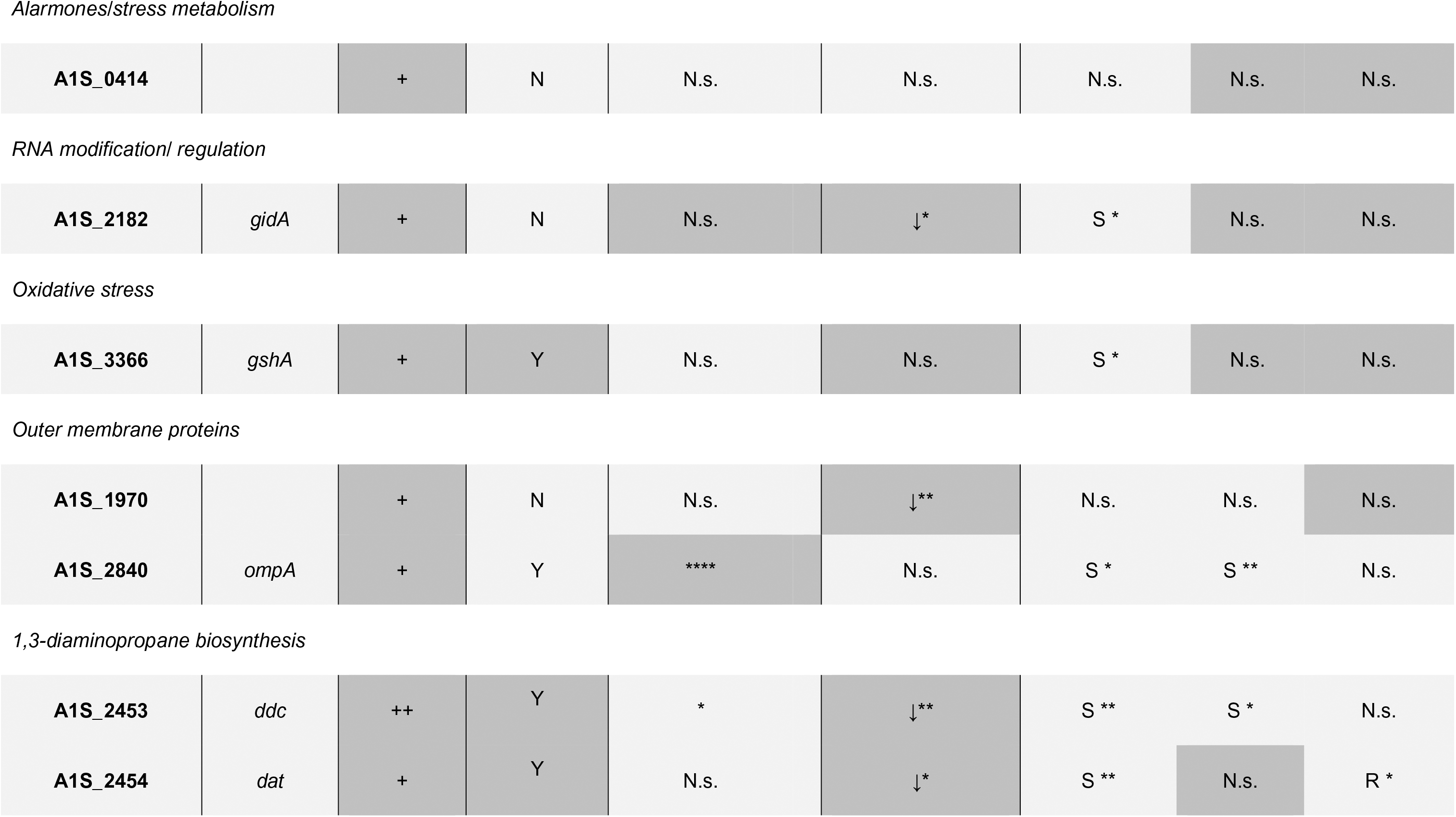
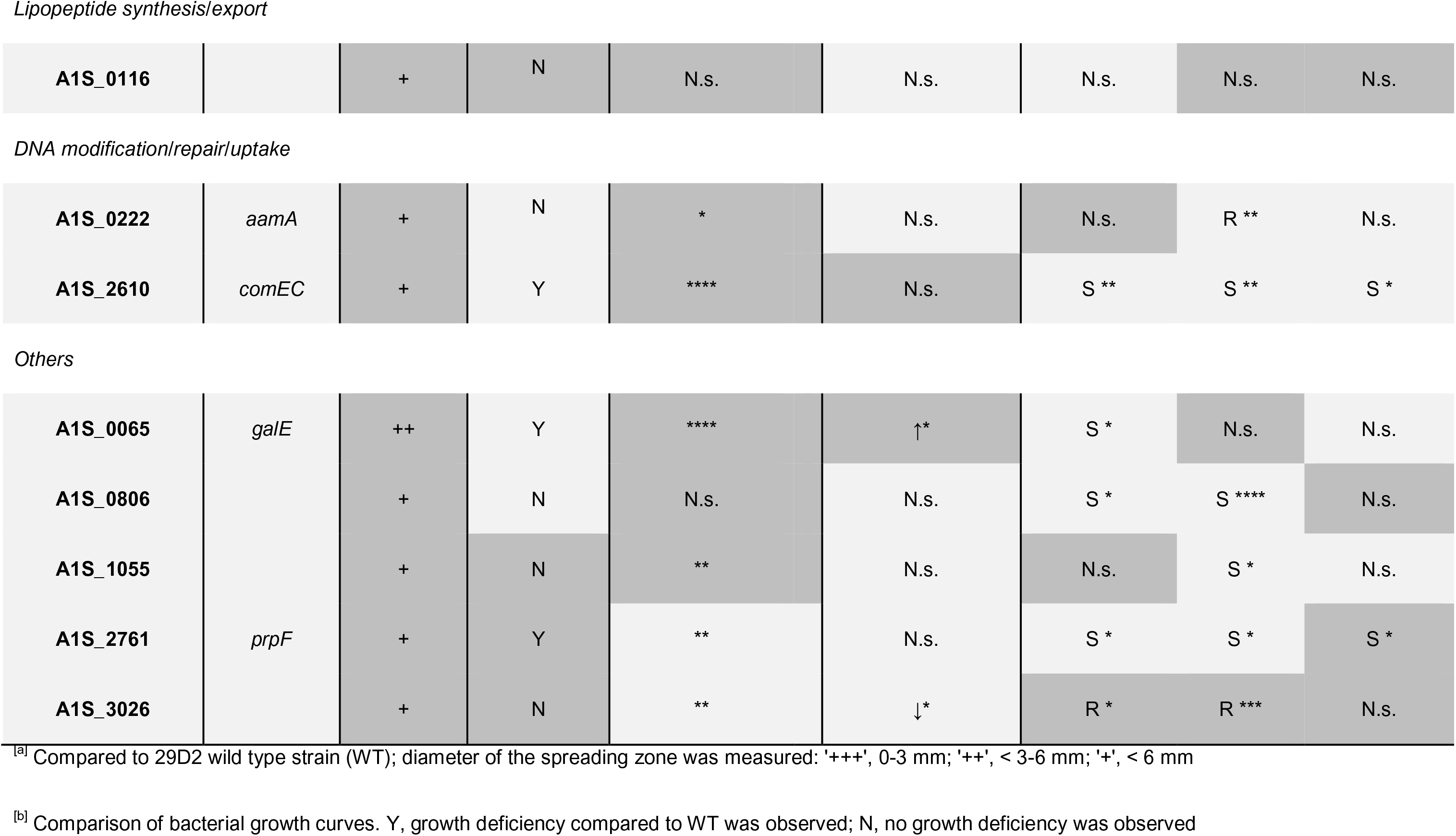
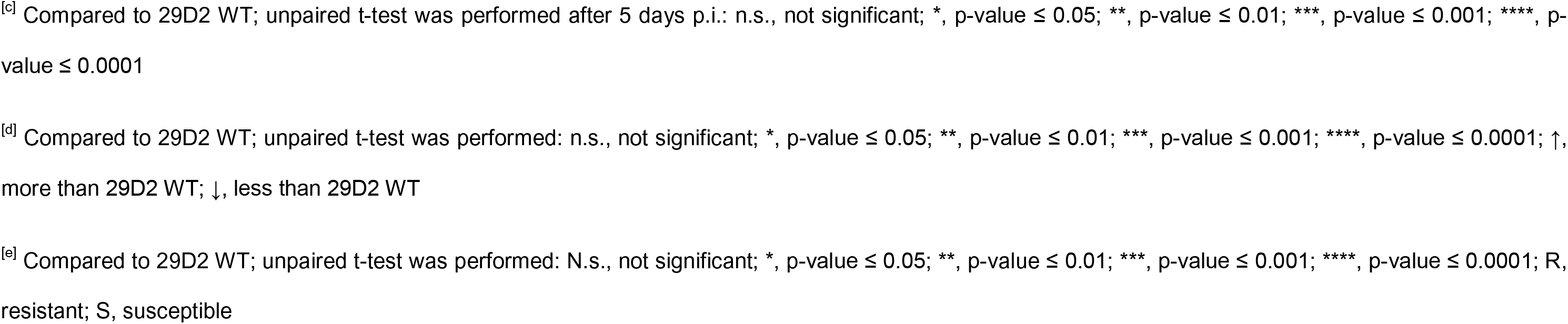
Summary of experimental results on genes involved in surface-associated motility in *A. baumannii* 29D2. A dark grey background indicates concordance to results obtained for strain ATCC 17978

| Locus tag in ATCC 17978 | Annotation/gene name | Motility deficiency<sup>[a]</sup> | Growth deficiency<sup>[b]</sup> | Attenuation in *Galleria mellonella* infection<sup>[c]</sup> | Pellicle biofilm formation<sup>[d]</sup> | MIC values<sup>[e]</sup> |  |  |
| --- | --- | --- | --- | --- | --- | --- | --- | --- |
|  |  |  |  |  |  | Ampicillin | Imipenem | Tetracycline |
| *Purine/pyrimidine/folate biosynthesis* |  |  |  |  |  |  |  |  |
| A1S_2187 | *purH* | ++ | Y | * | ↑* | S * | N.s. | N.s. |
| A1S_2251 | *purF* | ++ | Y | N.s. | N.s. | S ** | S * | N.s. |
| A1S_2605 | *purM* | + | Y | N.s. | N.s. | S * | N.s. | N.s. |
| A1S_2964 | *purE* | + | Y | N.s. | N.s. | S ** | N.s. | N.s. |
| A1S_1566 |  | + | Y | *** | N.s. | S * | N.s. | S * |
| A1S_2687 | *carB* | + | Y | **** | ↓** | S ** | S ** | N.s. |
| *Alarmones/stress metabolism* |  |  |  |  |  |  |  |  |
| **A1S_0414** |  | + | N | N.s. | N.s. | N.s. | N.s. | N.s. |
| *RNA modification/ regulation* |  |  |  |  |  |  |  |  |
| **A1S_2182** | *gidA* | + | N | N.s. | ↓* | S * | N.s. | N.s. |
| *Oxidative stress* |  |  |  |  |  |  |  |  |
| **A1S_3366** | *gshA* | + | Y | N.s. | N.s. | S * | N.s. | N.s. |
| *Outer membrane proteins* |  |  |  |  |  |  |  |  |
| **A1S_1970** |  | + | N | N.s. | ↓** | N.s. | N.s. | N.s. |
| **A1S_2840** | *ompA* | + | Y | **** | N.s. | S * | S ** | N.s. |

*1,3-diaminopropane biosynthesis*

|  |  |  |  |  |  |  |  |  |
| --- | --- | --- | --- | --- | --- | --- | --- | --- |
| **A1S_2453** | *ddc* | ++ | Y | * | ↓** | S ** | S * | N.s. |
| **A1S_2454** | *dat* | + | Y | N.s. | ↓* | S ** | N.s. | R * |
| *Lipopeptide synthesis/export* |  |  |  |  |  |  |  |  |
| **A1S_0116** |  | + | N | N.s. | N.s. | N.s. | N.s. | N.s. |
| *DNA modification/repair/uptake* |  |  |  |  |  |  |  |  |
| **A1S_0222** | *aamA* | + | N | * | N.s. | N.s. | R ** | N.s. |
| **A1S_2610** | *comEC* | + | Y | **** | N.s. | S ** | S ** | S * |
| *Others* |  |  |  |  |  |  |  |  |
| **A1S_0065** | *galE* | ++ | Y | **** | ↑* | S * | N.s. | N.s. |
| **A1S_0806** |  | + | N | N.s. | N.s. | S * | S **** | N.s. |
| **A1S_1055** |  | + | N | ** | N.s. | N.s. | S * | N.s. |
| **A1S_2761** | *prpF* | + | Y | ** | N.s. | S * | S * | S * |
| **A1S_3026** |  | + | N | ** | ↓* | R * | R *** | N.s. |

<sup>[a]</sup> Compared to 29D2 wild type strain (WT); diameter of the spreading zone was measured: '+++', 0-3 mm; '++', < 3-6 mm; '+', < 6 mm
<sup>[b]</sup> Comparison of bacterial growth curves. Y, growth deficiency compared to WT was observed; N, no growth deficiency was observed
<sup>[c]</sup> Compared to 29D2 WT; unpaired t-test was performed after 5 days p.i.: n.s., not significant; \*, p-value $\leq 0.05$ ; \*\*, p-value $\leq 0.01$ ; \*\*\*, p-value $\leq 0.001$ ; \*\*\*\*, p-value $\leq 0.0001$
<sup>[d]</sup> Compared to 29D2 WT; unpaired t-test was performed: n.s., not significant; \*, p-value $\leq 0.05$ ; \*\*, p-value $\leq 0.01$ ; \*\*\*, p-value $\leq 0.001$ ; \*\*\*\*, p-value $\leq 0.0001$ ; ↑, more than 29D2 WT; ↓, less than 29D2 WT
<sup>[e]</sup> Compared to 29D2 WT; unpaired t-test was performed: N.s., not significant; \*, p-value $\leq 0.05$ ; \*\*, p-value $\leq 0.01$ ; \*\*\*, p-value $\leq 0.001$ ; \*\*\*\*, p-value $\leq 0.0001$ ; R, resistant; S, susceptible

**Fig. 3.**
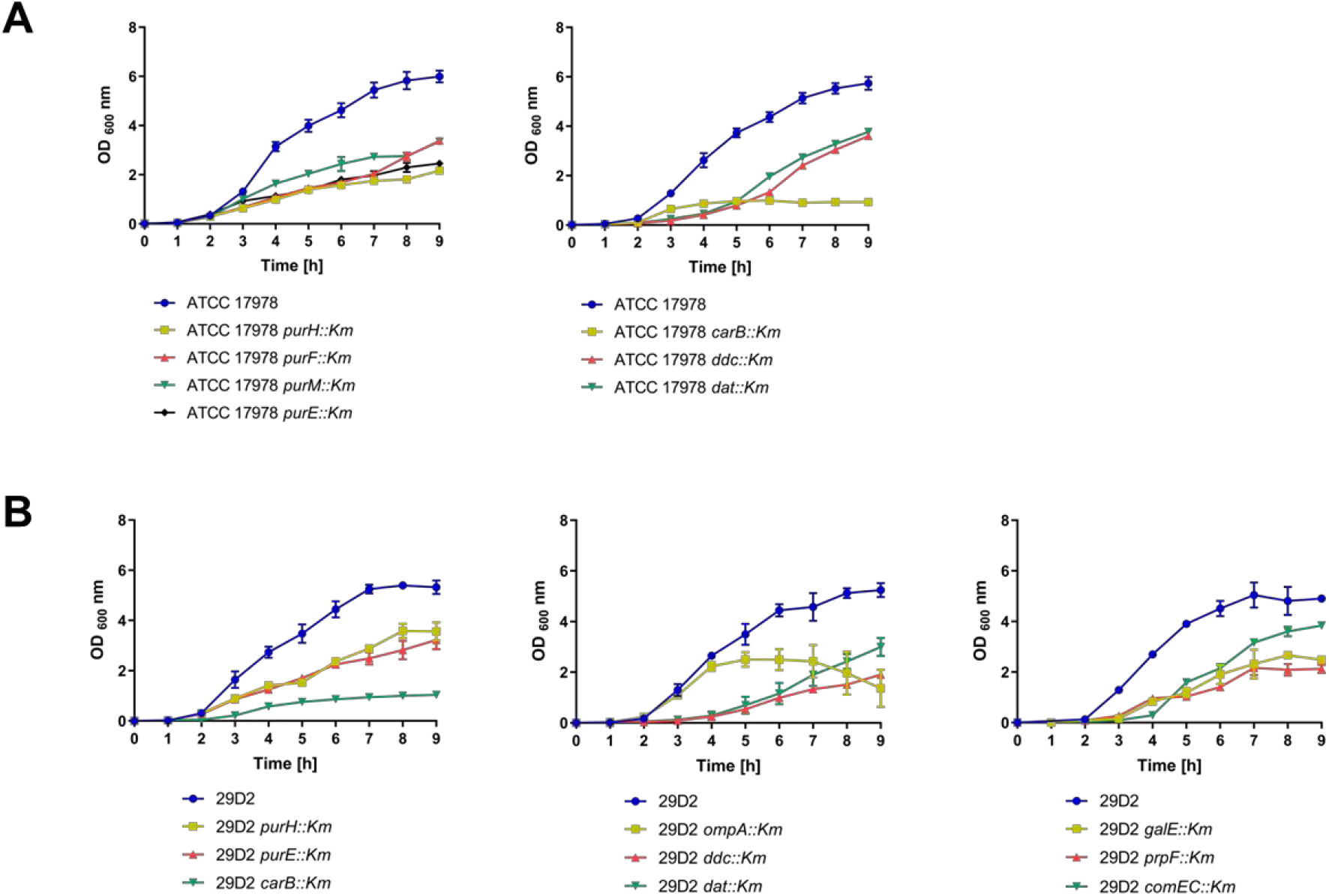
Growth deficiency of mutant strains from ATCC 17978 (A) and 29D2 (B) mutant libraries. OD-adjusted bacterial cultures were grown for 9 h at 37°C under constant shaking. Every hour cultures were measured at an OD of 600 nm. For each strain data obtained from 3 independent cultures grown on the same day were averaged and represented by the mean ± SD. Growth defects compared to wildtype ATCC 17978 were observed for mutants that are involved in purine, pyrimidine, and diaminopropane biosynthesis (**A**). In strain 29D2, growth defects were observed for mutants involved in purine/pyrimidine/folate and diaminopropane biosynthesis, and for mutants *galE::Km*, *ompA::Km,* and *prpF::Km* (**B**). See Supplementary Figs. S4 and S5 for growth curves of all strains described in this study.

In summary, we found that genes involved in purine/pyrimidine and diaminopropane biosynthesis, oxidative stress, and propionate catabolism were crucial for growth of ATCC 17978 and 29D2 in LB medium.

### *G. mellonella* caterpillar infection

To gain insight into a possible correlation between motility and virulence we made use of the *G. mellonella* infection model. Caterpillars were infected with 3 x 10^5^ CFU of different *A. baumannii* strains and the death of larvae was monitored over a time period of 5 days. *G. mellonella* infection of ATCC 17978 wildtype and mutant strains is shown in supplementary Fig. S6 and levels of significance for 5 days post-infection are presented in Table 1. A detailed listing of p-values for every monitored timepoint is provided in supplementary Table S6. After 24 h post-infection with the 17978 wildtype strain about 60% of larvae were dead. This number increased to over 80% of dead larvae after 5 days post-infection. 15 of 30 tested mutant strains displayed a significant attenuation in *G. mellonella* infection (Table 1). Another 4 mutant strains (*purE::Km*, *1624::Km*, *rpmG::Km*, and *ddc::Km*) showed some attenuation but this was not significant. The remaining 11 mutant strains did not display attenuation (Suppl. Fig. S6 and Table 1). Most pronounced attenuation was observed in strains *carB::Km*, *metG::Km*, *ompA::Km,* and *galE::Km* (Fig. 4A). These results suggest an important role for these genes in *A. baumannii* virulence. However, to exclude the possibility that attenuation could be due to decreased planktonic growth, we compared the caterpillar infection results to our bacterial growth data (Fig. 3, Suppl. Fig. S4, and Fig. S5). Among the above mentioned mutants, only the *galE*::*Km* mutant was not significantly affected in growth. Overall, we found that for 11 of 15 significantly attenuated mutant strains the caterpillar infection data could possibly be influenced by decreased growth rates (Table 1 and Fig. 3).

**Fig. 4.**
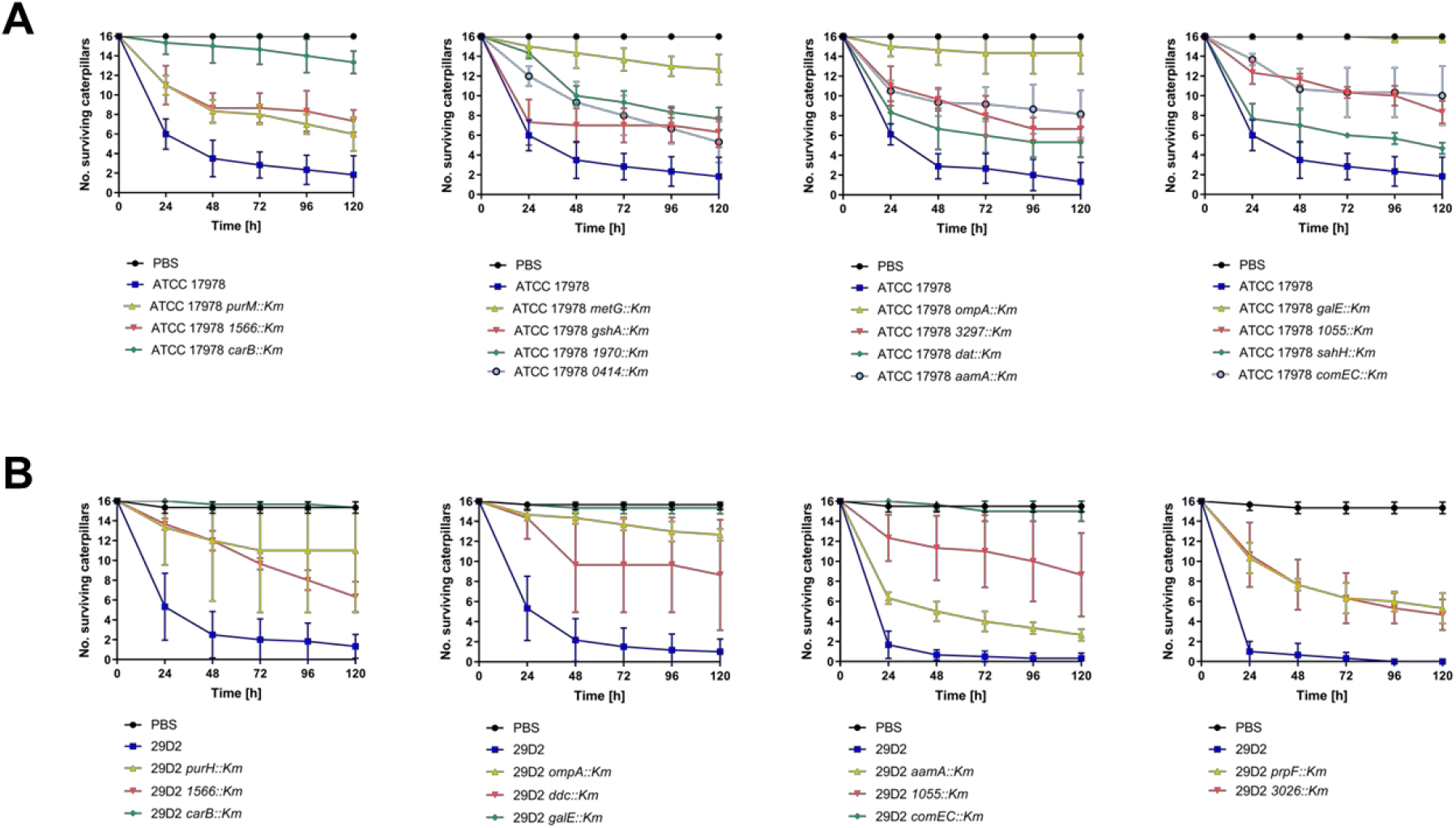
Attenuation of *A. baumannii* ATCC 17978 mutants (A) and 29D2 mutants (B) in the *G. mellonella* caterpillar infection model. Caterpillars were infected with 3 x 10^5^ CFU of *A. baumannii* strains as indicated. Sterile PBS (black lines) was used as a control. Three independent experiments were performed with groups of 16 caterpillars for every bacterial strain and control. Data obtained from 3 independent experiments were averaged and represented by the mean ± SD. In strain ATCC 17978, 15 of 30 mutants showed a significant attenuation at 5 days post-infection (see Table 1 for p-values) compared to the wildtype strain (**A**). In strain 29D2, 11 of 21 mutants were attenuated (see Table 2 for p-values) in the *G. mellonella* infection model (**B**). See Supplementary Figs. S6 and S7 for infection data of all strains described in this study.

The *G. mellonella* infection with 29D2 wildtype and mutant strains data is shown in supplementary Fig. S7 and significance levels (for 5 days post-infection) are shown in Table 2. A detailed listing of p-values for every monitored timepoint is provided in supplementary Table S7. 11 of 21 29D2 mutants were significantly attenuated in the *G. mellonella* infection model (Suppl. Fig S7 and Table 2). Within this group the most pronounced attenuation was observed in strains *carB::Km*, *ompA::Km*, *galE::Km,* and *comEC::Km* (Fig. 4B). The mutant strains *purE::Km*, *gidA::Km,* and *0806::Km* showed some attenuation at 5 days post-infection but p-values ranged between 0.060 – 0.067 (Suppl. Table S7). Interestingly, 8 of 11 significantly attenuated mutant strains (*purH::Km*, *1566::Km*, *carB::Km*, *ompA::Km*, *ddc::Km*, *comEC::Km*, *galE::Km*, and *2761::Km*) manifested a growth deficiency compared to the parental strain (Suppl. Fig. S5 and Table 2).

In summary, concordant infection traits were observed for 12 mutants of both strains including mutants affected in purine/pyrimidine/folate biosynthesis. Among these 12 strains, most significant attenuation was observed for *carB::Km*, *ompA::Km,* and *galE::Km*.

As a control, the CFUs were determined from the OD-adjusted bacterial cultures used for the infection experiments. To this end, OD-adjusted cultures were serially diluted and plated on nutrient agar. Colonies were counted after incubation for 18 h at 37°C. Interestingly, for both ATCC 17978 *ompA::Km* and 29D2 *ompA::Km* mutants we observed 1-2 log scale lower CFU numbers compared to the OD-adjusted suspension (data not shown). Sticking of cells to the tube wall during the dilution process was minimized by using low-binding tubes (Eppendorf). When growing both *ompA::Km* mutants on agar plates we observed a very sticky colony texture upon touching with a glass rod. Based on these findings we examined the cell morphology of *ompA::Km* mutant strains under the microscope. A distinct cell elongation or chain formation of both *ompA::Km* mutant strains compared to their parental strains was observed (Supplementary Fig. S8).

### MIC Determination

We aimed to elucidate the correlation between motility-deficient mutants and their sensitivity to the bactericidal antibiotics ampicillin and imipenem as well as to the bacteriostatic antibiotic tetracycline. For ATCC 17978, 18 of 30 mutants displayed a significant resistance to ampicillin compared to the parental strain. The highest MIC values were obtained in mutant strains *0414::Km* (4-fold increase compared to the parental strain)*, 3026::Km* (4-fold increase), and *1566::Km* (3.7-fold increase). The only mutant strain which showed decreased resistance (0.7-fold decrease) to ampicillin was *aamA::Km* (Table 3). By contrast, a significantly increased sensitivity to imipenem was observed in six (*purE::Km*, *carB::Km*, *1624::Km, rpmG::Km*, *1970::Km*, and *aamA::Km*) of the tested strains (Table 1). Furthermore, a significantly increased resistance to imipenem was observed in 4 of the tested mutant strains (*ompA::Km*, *3297::Km*, *0806::Km*, and *3026::Km*). For tetracycline, we found 13 of 30 mutants to be significantly more sensitive compared to the parental strain. Only 2 of 30 mutant strains, *purF::Km* and *galE::Km*, displayed significantly increased resistance to tetracycline (Table 1). Next, we analyzed all 29D2 mutant strains with respect to the MIC values for ampicillin, imipenem, and tetracycline. A significantly increased sensitivity to ampicillin was observed in 15 of 21 tested mutant strains compared to wildtype (Table 2). The only strain with significantly increased resistance to ampicillin was *3026::Km*, with a 1.7–fold increased MIC value (Table 3). Another mutant strain with a 1.5-fold increased ampicillin MIC value, although not significant, was *aamA::Km*. Similar effects were observed for imipenem (Table 3). Here, strains *3026::Km* and *aamA::Km* displayed significant resistance compared to the parental strain (Table 2). Increased sensitivity to imipenem was observed in 8 of 21 tested mutants. For the MIC values of tetracycline, we found the 3 mutant strains *1566::Km* (3.6-fold decrease), *comEC::Km* (5.4-fold decrease), and *prpF::Km* (6.5-fold decrease) to be significantly more susceptible compared to the parental strain (Table 3). Only one mutant, *dat::Km*, was significantly more resistant to tetracycline with a 1.8-fold increase (Table 3).

**Table 3.**
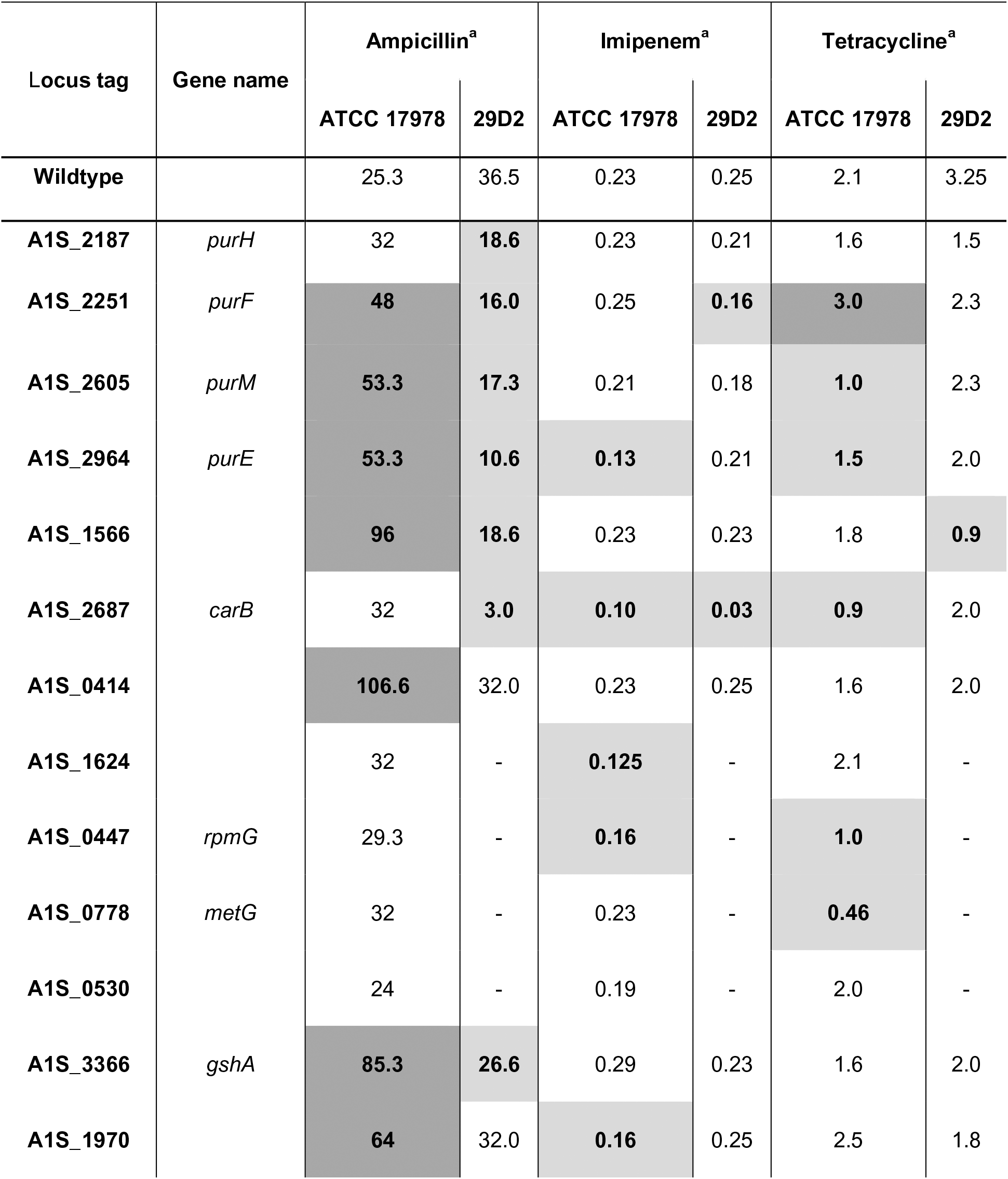
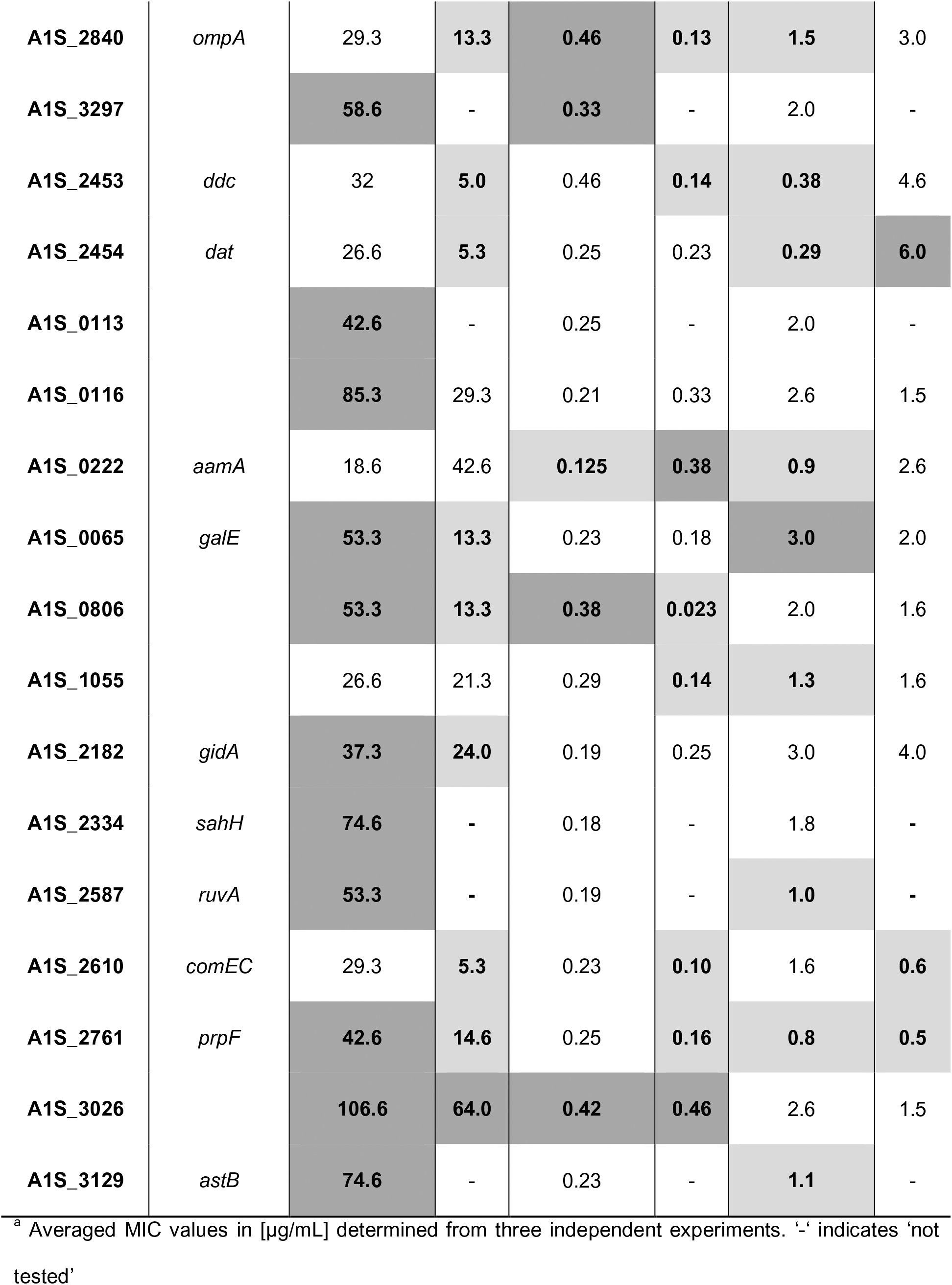
Minimal inhibitory concentration (MIC) of ampicillin, tetracycline and imipenem determined from ATCC 17978 wildtype/mutants and 29D2 wildtype/mutants. A dark grey background indicates that MIC values of mutant strains are significantly elevated compared to the wildtype while a light grey background indicates increased susceptibility

| Locus tag | Gene name | Ampicillin<sup>a</sup> |  | Imipenem<sup>a</sup> |  | Tetracycline<sup>a</sup> |  |
| --- | --- | --- | --- | --- | --- | --- | --- |
|  |  | ATCC 17978 | 29D2 | ATCC 17978 | 29D2 | ATCC 17978 | 29D2 |
| **Wildtype** |  | 25.3 | 36.5 | 0.23 | 0.25 | 2.1 | 3.25 |
| **A1S_2187** | *purH* | 32 | **18.6** | 0.23 | 0.21 | 1.6 | 1.5 |
| **A1S_2251** | *purF* | **48** | **16.0** | 0.25 | **0.16** | **3.0** | 2.3 |
| **A1S_2605** | *purM* | **53.3** | **17.3** | 0.21 | 0.18 | **1.0** | 2.3 |
| **A1S_2964** | *purE* | **53.3** | **10.6** | **0.13** | 0.21 | **1.5** | 2.0 |
| **A1S_1566** |  | **96** | **18.6** | 0.23 | 0.23 | 1.8 | **0.9** |
| **A1S_2687** | *carB* | 32 | **3.0** | **0.10** | **0.03** | **0.9** | 2.0 |
| **A1S_0414** |  | **106.6** | 32.0 | 0.23 | 0.25 | 1.6 | 2.0 |
| **A1S_1624** |  | 32 | - | **0.125** | - | 2.1 | - |
| **A1S_0447** | *rpmG* | 29.3 | - | **0.16** | - | **1.0** | - |
| **A1S_0778** | *metG* | 32 | - | 0.23 | - | **0.46** | - |
| **A1S_0530** |  | 24 | - | 0.19 | - | 2.0 | - |
| **A1S_3366** | *gshA* | **85.3** | **26.6** | 0.29 | 0.23 | 1.6 | 2.0 |
| **A1S_1970** |  | **64** | 32.0 | **0.16** | 0.25 | 2.5 | 1.8 |
| **A1S_2840** | *ompA* | 29.3 | **13.3** | **0.46** | **0.13** | **1.5** | 3.0 |
| **A1S_3297** |  | **58.6** | - | **0.33** | - | 2.0 | - |
| **A1S_2453** | *ddc* | 32 | **5.0** | 0.46 | **0.14** | **0.38** | 4.6 |
| **A1S_2454** | *dat* | 26.6 | **5.3** | 0.25 | 0.23 | **0.29** | **6.0** |
| **A1S_0113** |  | **42.6** | - | 0.25 | - | 2.0 | - |
| **A1S_0116** |  | **85.3** | 29.3 | 0.21 | 0.33 | 2.6 | 1.5 |
| **A1S_0222** | *aamA* | 18.6 | 42.6 | **0.125** | **0.38** | **0.9** | 2.6 |
| **A1S_0065** | *galE* | **53.3** | **13.3** | 0.23 | 0.18 | **3.0** | 2.0 |
| **A1S_0806** |  | **53.3** | **13.3** | **0.38** | **0.023** | 2.0 | 1.6 |
| **A1S_1055** |  | 26.6 | 21.3 | 0.29 | **0.14** | **1.3** | 1.6 |
| **A1S_2182** | *gidA* | **37.3** | **24.0** | 0.19 | 0.25 | 3.0 | 4.0 |
| **A1S_2334** | *sahH* | **74.6** | - | 0.18 | - | 1.8 | - |
| **A1S_2587** | *ruvA* | **53.3** | - | 0.19 | - | **1.0** | - |
| **A1S_2610** | *comEC* | 29.3 | **5.3** | 0.23 | **0.10** | 1.6 | **0.6** |
| **A1S_2761** | *prpF* | **42.6** | **14.6** | 0.25 | **0.16** | **0.8** | **0.5** |
| **A1S_3026** |  | **106.6** | **64.0** | **0.42** | **0.46** | 2.6 | 1.5 |
| **A1S_3129** | *astB* | **74.6** | - | 0.23 | - | **1.1** | - |

<sup>a</sup> Averaged MIC values in [µg/mL] determined from three independent experiments. '-' indicates 'not tested'

**Table 4.**
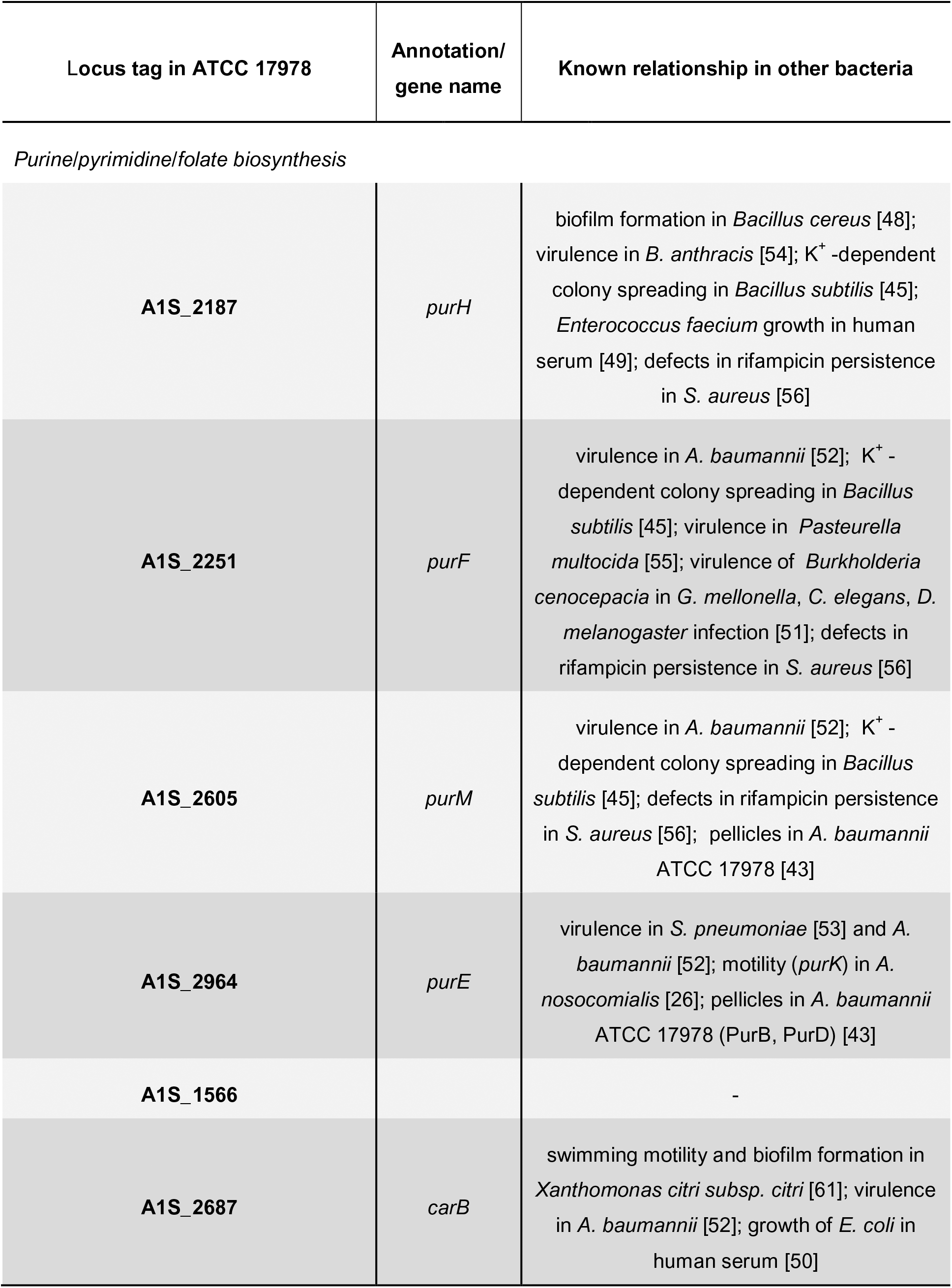
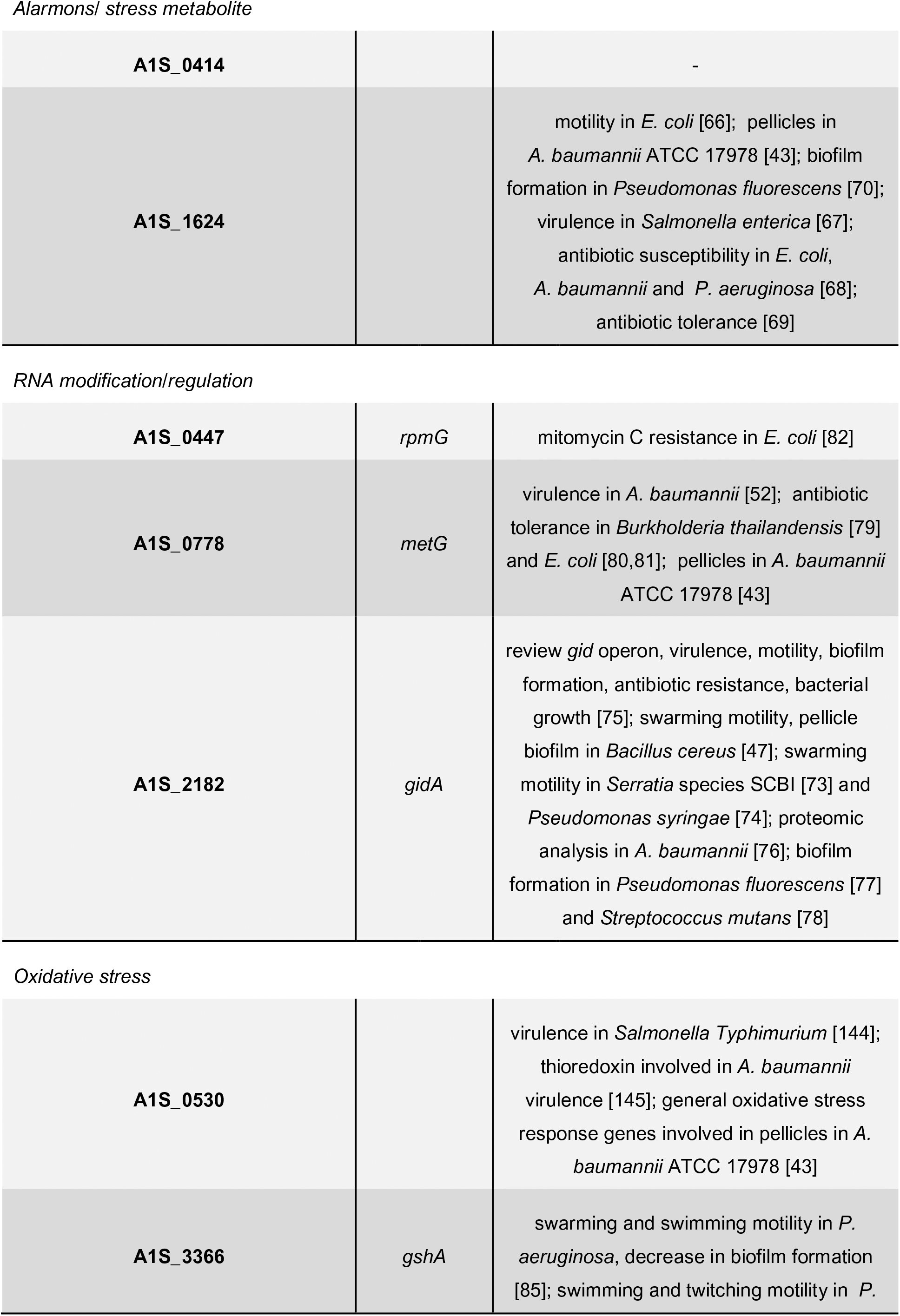
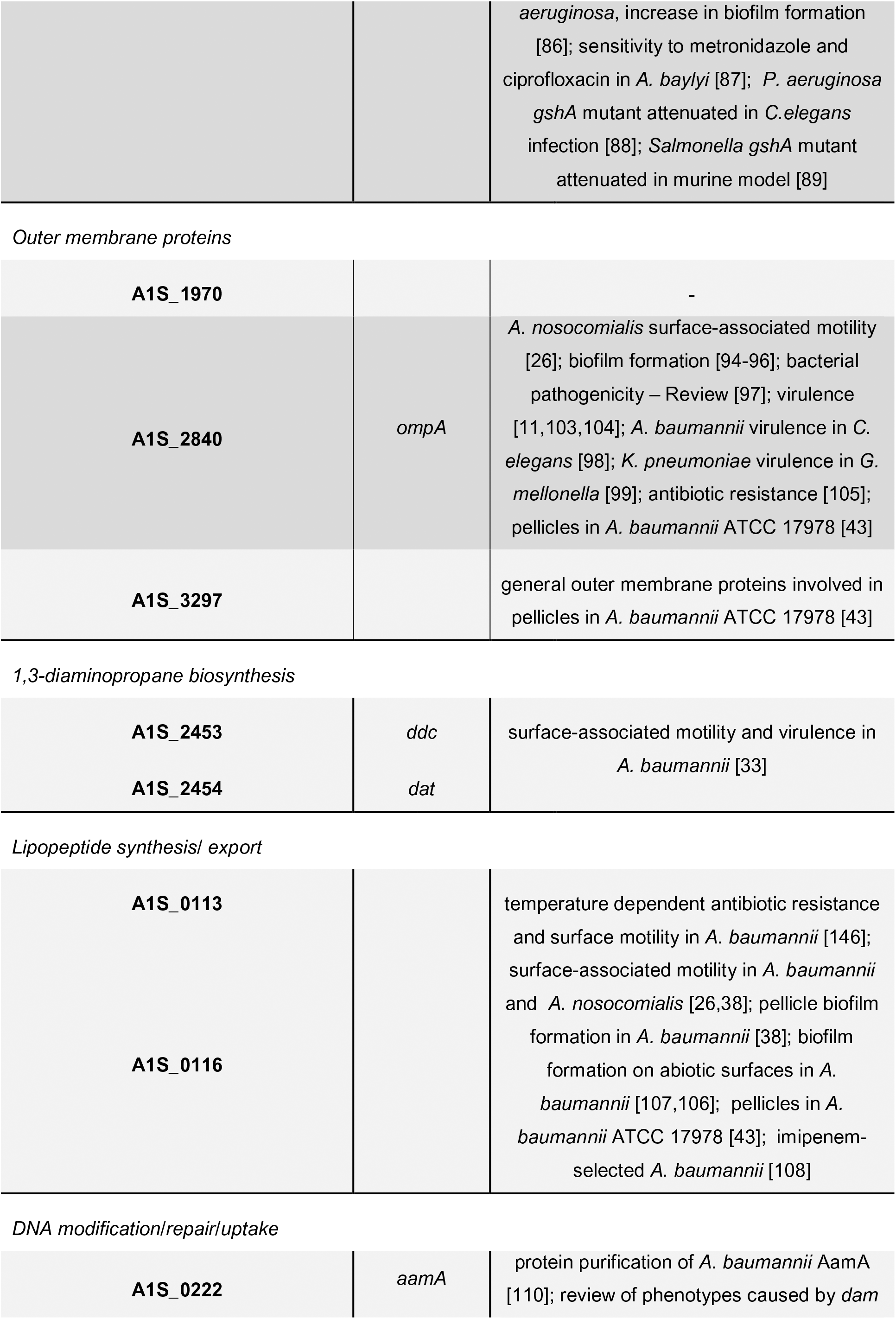
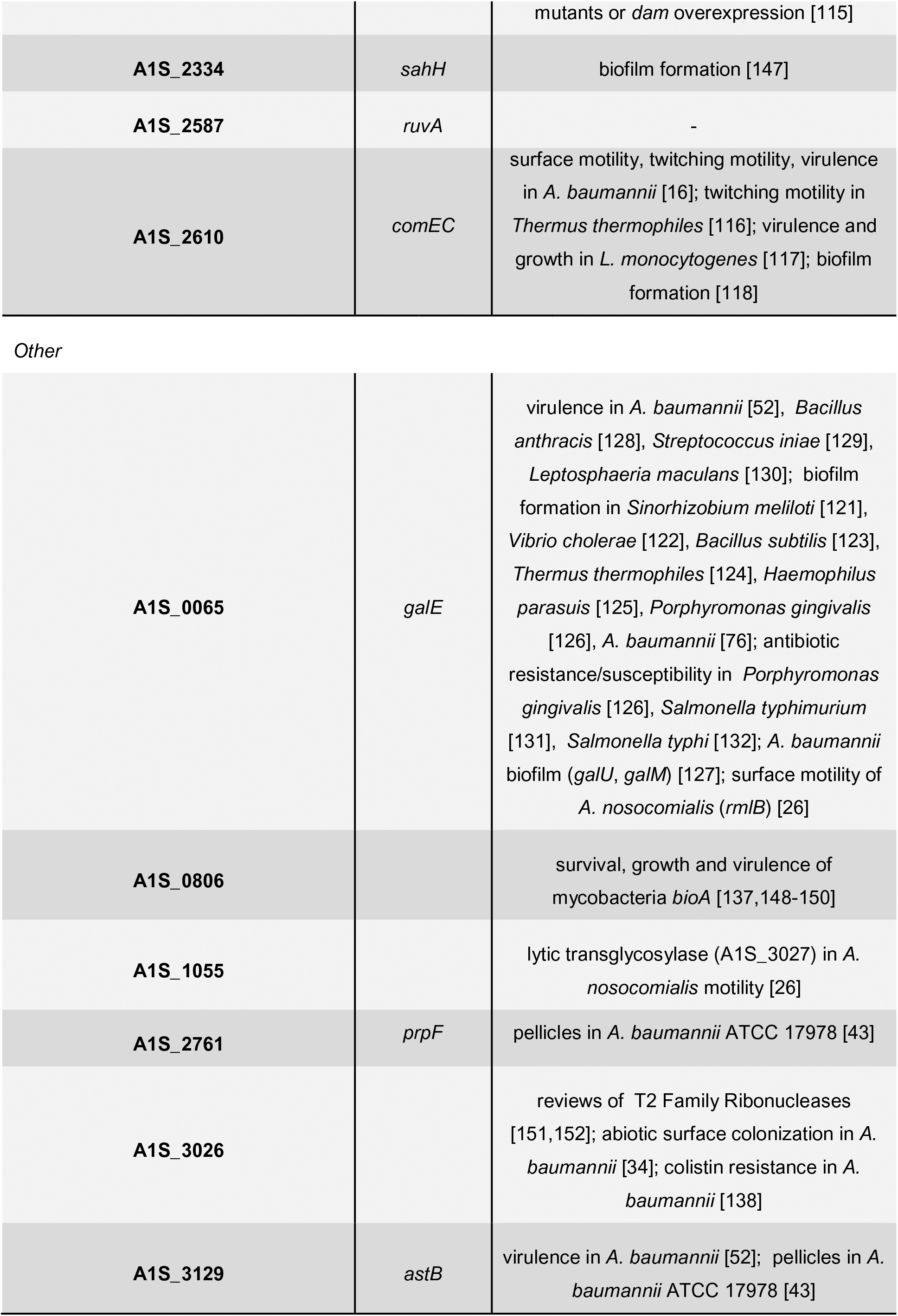
Links between the genes identified in this study and the literature.

In conclusion, mutants from the 29D2 background predominantly showed increased sensitivity to all tested antibiotics. By contrast, many mutants of ATCC 17978 showed increased resistance to ampicillin, but increased sensitivity to imipenem and tetracycline.

## Discussion

Recently, *A. baumannii* was demonstrated to exhibit motility on semi-dry plates, with agar concentrations between 0.2-0.4%, and motility was dependent on the type of agar that was used. Bacterial surface spreading was shown not to depend on type IV pili [26]. Here, we characterized 30 genes involved in *A. baumannii* surface-associated motility with respect to bacterial growth, pellicle biofilm formation, virulence, and antibiotic resistance. We discuss motility-deficient mutants with regards to their known/putative gene function in the bacterial cell (Fig. 5).

**Fig. 5.**
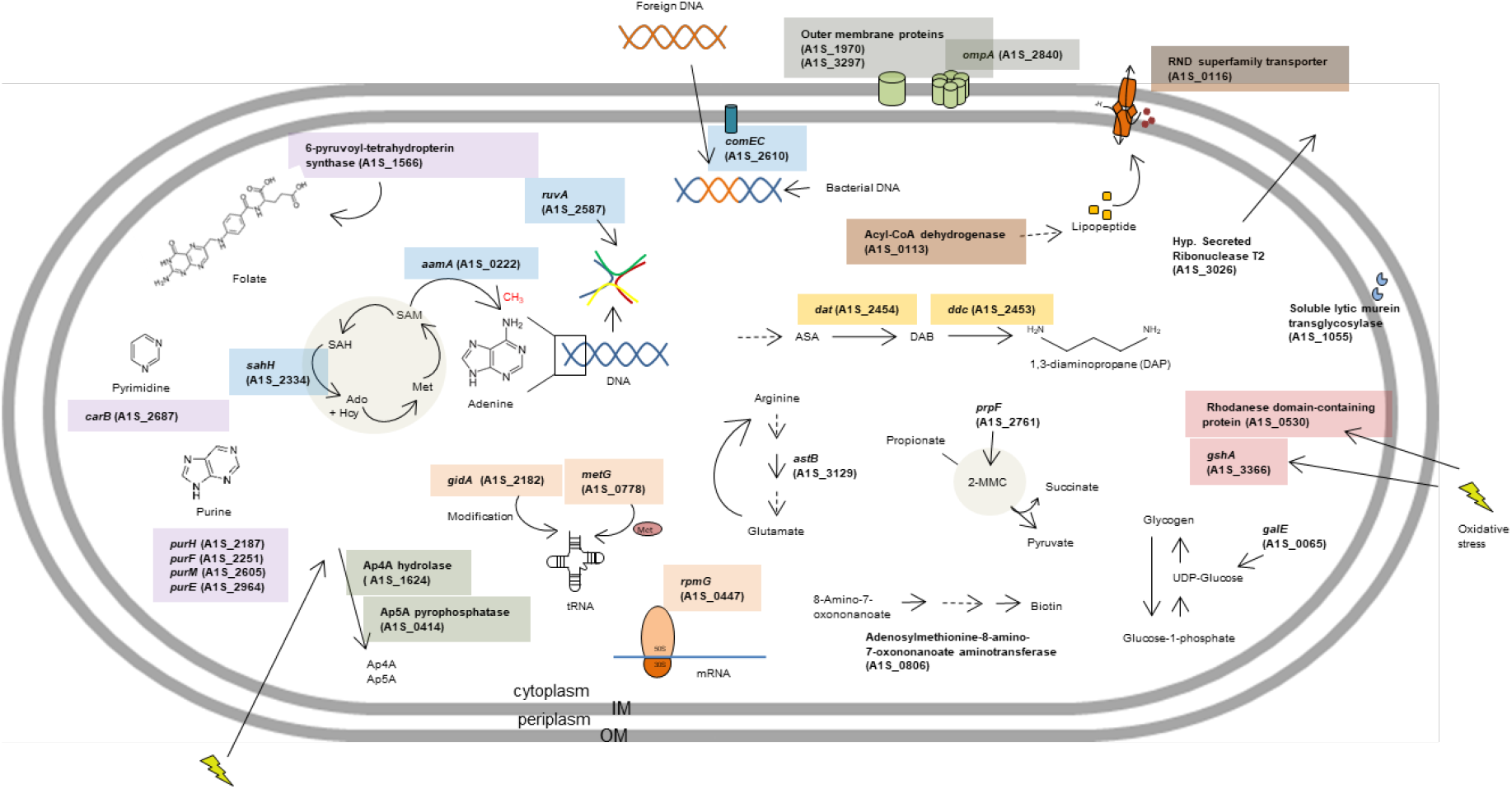
Genes inactivated in *A. baumannii* ATCC 17978 mutants with a surface-associated motility defect and their known/predicted/putative function in the bacterial cell. A common color code indicates that mutants belong to the same functional, processual, and/or structural category. OM, outer membrane; IM, inner membrane; Ap4A, diadenosine tetraphosphate; Ap5A, diadenosine pentaphosphate; SAM, S-adenosyl-L-methionine; SAH, S-adenosylhomocysteine; Ado, adenosine; Hcy, homocysteine; Met, methionine; ASA, L-aspartate 4-semialdehyde; DAB, L-2,4-diaminobutanoate; 2-MMC, 2-methylcitric acid cycle.

### Genes involved in purine/pyrimidine/folate biosynthesis

In our study we identified 4 proteins involved in purine (*pur*) biosynthesis to be essential for *A. baumannii* surface-associated motility: PurH, PurF, PurM, and PurE (Tables 1 and 2). The decrease in surface motility of these mutants ranged from a 25-fold decrease in ATCC 17978 to a 2.1-fold decrease in 29D2 compared to their parental strains (Suppl. Table S3). In *A. nosocomialis* strain M2, EZ::Tn insertion in gene *purK* (*a1s_2963*) has been previously described to result in a 70% reduction in surface motility compared to the parental strain [26]. Mutations in the genes *purD*, *purF*, *purH*, *purL*, and *purM* abolished K^+^-dependent colony spreading in *Bacillus subtilis* [46]. The *pur* genes were also demonstrated to be essential for biofilm formation in *Bacillus cereus* (*purH* and *purD* [47]; *purA* [48]; *purA*, *purC*, and *purL* [49]). Interestingly, our study revealed no defective role of *pur* genes in pellicle biofilm formation. In contrast, mutations *purH::Km*, *purM::Km*, and *purE::Km* in 17978 and *purH::Km* in 29D2 produced significantly more pellicle biomass than their parental strains (Tables 1 and 2). A pellicle proteome study in 17978 showed that the *pur* proteins were differentially expressed/accumulated under planktonic (PurH, PurF, and PurA), 1-day pellicle (PurM and PurB), and 4-day pellicle (PurD) growth conditions [44].

In addition to the motility deficiency, we found all tested *pur* mutants to display bacterial growth defects in LB media (Suppl. Figs. S4 and S5). For various bacteria, *pur* genes were identified to be required for bacterial growth in human serum including *Enterococcus faecium* (*purL*, *purH*, and *purD*) [50], *E. coli* (*purA*, *C*, *D*, *E*, *F*, *H*, *K*, *L*, and *M*), *S. enterica* (*purA*, *B*, *C*, *D*, *E*, *F*, *G*, and *H*), and *B. anthracis* (*purE* and *purK*) [51]. Due to the fact that all *pur* mutants displayed bacterial growth defects we expected these mutants to be attenuated in the *G. mellonella* infection, but we only found the 2 mutants ATCC 17978 *purM::Km* and 29D2 *purH::Km* had significant attenuation (Fig. 4). Purine biosynthesis mutants (*purF*, *purD*, and *purL*) in *Burkholderia cenocepacia* were also found to be attenuated in the *G. mellonella* infection model as well as in *C. elegans* and *D. melanogaster* infections [52]. *De novo* purine biosynthesis has also been shown to be required for virulence in ATCC 17978 (*purF*, *purD*, *purN*, *purL*, *purM*, *purK*, *purE*, *purC*, *purP*, and *purO*) in the mouse lung [53], and in several other bacteria such as *Streptococcus pneumoniae* (*purE*, *purK*, *purC*, and *purL*) [54], *Bacillus anthracis* (*purH*) [55], and *Pasteurella multocida* (*purF*) [56].

The *pur* mutants were tested for antibiotic sensitivity/resistance against ampicillin, imipenem, and tetracycline. For ampicillin MIC’s we obtained very contrary results. The *pur* mutants of 29D2 showed increased sensitivity to ampicillin, but, except for *purH::Km,* the *pur* mutants of ATCC 17978 were significantly more resistant compared to their parental strain (Table 3). In *S. aureus* defective rifampicin persistence was shown for *purB*, *purF*, *purH,* and *purM* [57].

The *A. baumannii* gene *a1s_2687* encodes the large subunit (*carB*) of carbamoylphosphate synthase and is arranged with the small subunit (*carA*) in the *carAB* operon, which is required for the *de novo* synthesis of arginine and pyrimidines (reviewed in [58]), and in turn pyrimidines are known to be involved in biofilm formation in *P. aeruginosa* [59] and *E. coli* [60]. In *A. baumannii*, inactivation of *carB* caused a significantly decreased persistence in a mouse pneumonia model [53]. The contribution of *carB* to *A. baumannii* virulence was confirmed by our results, showing significant attenuation in both ATCC 17978 *carB::Km* and 29D2 *carB::Km* mutant strains (Fig. 4). Additionally, in a *P. aeruginosa* competition study against *B. cepacia*, *K. pneumoniae*, and *S. aureus*, the *carB* gene and hence uracil/pyrimidine biosynthesis was identified to be essential [61]. Inactivation of *carB* in ATCC 17978 and 29D2 resulted in the greatest phenotypic alterations in planktonic growth, pellicle biofilm formation, and *G. mellonella* caterpillar infection of all tested mutants (Tables 1 and 2). Interestingly, similar observations were also made for the gammaproteobacterium *Xanthomonas citri subsp. citri*. In that study, the knockout of *carB* resulted in a 70% decrease in biofilm formation, an extensive reduction in swimming motility, and alterations in bacterial growth [62]. CarB was also found to be required for growth of *E. coli* in human serum [51]. A motility-deficient phenotype was identified for the gene *a1s_1566* (putative 6-pyruvoyl-tetrahydropterin synthase), involved in folate biosynthesis and thus crucial for biosynthesis of purines and deoxythymidine monophosphate (Fig. 1). Here we observed an involvement in virulence, bacterial growth, and pellicle biofilm formation (Tables 1 and 2). Taken together, these findings suggest that purine and pyrimidine genes contribute to important bacterial processes like motility, bacterial growth, pellicle biofilm formation, and virulence not only in *Acinetobacter* but also in well studied genera like *Bacillus* and *Salmonella*.

### Genes involved in alarmone/damage metabolism

The *A. baumannii* genes *a1s_0414* and *a1s_1624* encode for an Ap5A pyrophosphatase and an Ap4A hydrolase (ApaH-like), respectively, and are proposed to be involved in depletion of putative alarmones/signaling molecules [63, 64] and/or damage metabolites [65, 66]. Recent work suggests that dinucleoside polyphosphates can be used by RNA polymerases to initiate transcription and to act as 5’-RNA caps that may stabilize RNA, while ApaA-like hydrolases are able to remove these caps [67]. The Ap4A hydrolase knockout mutant *a1s_1624::Km* seems to play a role in *A. baumannii* surface motility and planktonic growth (Table 1). An *E. coli* Ap4A hydrolase (*apaH*) knockout mutant was previously associated with decreased motility [68]. In *Salmonella enterica* adhesion and invasion capacity into epithelial cells was reduced for the Δ*apaH* mutant [69]. Additionally, the *a1s_1624::Km* mutant exhibited increased sensitivity to imipenem (Table 1). Increased sensitivity of Δ*apaH* mutants against kanamycin and streptomycin was also shown for ATCC 17978, *E. coli,* and *P. aeruginosa* [70], and decreased sensitivity of Δ*apaH* mutants, in the form of persister cells, was found in *E. coli* [71]. Although we could not observe a significant impact of *a1s_1624::Km* on pellicle formation, others have shown A1S_1624 to be overproduced in pellicle cells in ATCC 17978 [44], and Ap4A metabolism impacts biofilm formation in *Pseudomonas fluorescens* [72]. The Ap5A pyrophosphatase knockout in ATCC 17978 resulted in a significant reduction in pellicle biofilm formation, significant attenuation of *G. mellonella* infection, and resistance against ampicillin (Table 1). Interestingly, in the corresponding *a1s_0414::Km* mutant of 29D2, only surface-associated motility was affected (Table 2).

In general, Ap4A and Ap5A are thought to be synthesized by aminoacyl-tRNA synthetases in the absence of tRNAs during amino acid activation. This process requires ATP and a cognate amino acid [73, 74]. Providing a possible link, we found a methionyl-tRNA synthetase in our surface motility-deficient library (discussed below).

### Genes involved in RNA modification/regulation

We found 3 genes involved in the regulation and/or modification of RNAs: *metG* (methionyl-tRNA synthetase, *a1s_0778*), *rpmG* (50S ribosomal protein L33, *a1s_0447*), and *gidA* (glucose-inhibited division protein A, a tRNA modification enzyme, *a1s_2182*). The deficiency in motility of *gidA* mutants has been described mainly for swarming motility in bacteria like *Bacillus cereus* [48], *Serratia* species SCBI [75], and *Pseudomonas syringae* [76]. In the present study, a *gidA* null allele in strains ATCC 17978 and 29D2 resulted in small decreases in their planktonic growth (Suppl. Fig. S4 and S5). Contrary results for Δ*gidA* bacterial growth has been reported (reviewed in [77]). Interestingly, proteomic analysis of *A. baumannii* planktonic and biofilm growth identified GidA only under biofilm growth conditions [78], while several studies reported the negative effect of *gidA* mutants on biofilm formation in different bacteria [79, 80]. In the present study we also saw a significant reduction of the pellicle-biofilm formation in both *gidA::Km* mutants (Fig. 2). An essential role of *gidA* in pellicle-biofilm formation was also shown in *Bacillus cereus* [48]. While several GidA-associated virulence effects have been reported (reviewed in [77]) we did not see significant attenuation in the *G. mellonella* infection model (Suppl. Figs. S6 and S7).

In contrast, the knockout of *metG* (*a1s_0778*) was associated with a significant attenuation in the *G. mellonella* infection model (Fig. 4). Similarly, involvement of *metG* in *A. baumannii* virulence was also shown in a mouse pneumonia model [53]. The *metG::Km* mutant revealed a significantly reduced ability to form pellicles (Fig. 2). Moreover, MetG was found to be more abundant in *A. baumannii* pellicle cells than in planktonic cells [44]. In our study we found the *metG::Km* mutant to be more sensitive to tetracycline (Table 1), which agrees with observations of amino acid substitutions of MetG associated with increased antibiotic tolerance in *Burkholderia thailandensis* [81] and *E. coli* [82, 83].

We observed increased sensitivity of the *rpmG::Km mutant* to imipenem and tetracycline (Table 1). This data is in line with a study which showed that a mitomycin C resistance phenotype was associated with RpmG overproduction in *E. coli* [84]

### Genes involved in oxidative stress

The ATCC 17978 gene *a1S_3366* is predicted to encode a gamma-glutamate-cysteine ligase (*gshA*) which is required to synthesize glutathione (GSH), an antioxidant molecule that protects cells against oxidative stress [85, 86]. Different studies observed a decrease in swarming [87], swimming [87, 88], and twitching motility [88] of the *P. aeruginosa* Δ*gshA* mutant compared to the parental strain. Contrary results were found for the ability *of P. aeruginosa* Δ*gshA* to form biofilms (increased in [88] and decreased in [87]). We did not find any changes in pellicle biofilm production compared to the parental strains for both of our *gshA* mutants (Fig. 2). In *Acinetobacter baylyi* the knockout of *gshA* increased sensitivity to metronidazole and ciprofloxacin [89]. We observed an enhanced sensitivity to ampicillin for the 29D2 *gshA::Km* mutant, but the ATCC 17978 *gshA::Km* mutant showed a resistant phenotype (Table 3). Attenuation in *G. mellonella* infection was observed for the ATCC 17978 *gshA* mutant strain, which agrees with other studies describing *gshA* mutants to be attenuated in *C. elegans* infection (*P. aeruginosa* [90]) and a murine infection model (*Salmonella* [91]).

The *A. baumannii* gene *a1s_0530* encodes for a rhodanese domain-containing protein, a putative sulfurtransferase, supposed to be involved in oxidative stress detoxification and sulfur metabolism [92–95]. The only knockout-related phenotype, besides surface-associated motility-deficiency, that we observed in ATCC 17978 was a significant increase in pellicle biofilm production (Fig. 2A). Finally, the involvement of oxidative stress response proteins in air-liquid pellicles has been described recently in a proteomic study of ATCC 17978 [44].

### Outer membrane proteins

The gene *a1s_3297* encodes a putative outer membrane protein and *a1s_1970* encodes a putative membrane-associated Zn-dependent protease (RseP). Here we show that both genes are involved in *A. baumannii* virulence, pellicle formation, and antimicrobial resistance (Tables 1 and 2).

We found OmpA to be involved in *A. baumannii* surface-associated motility, which has been described for the *A. nosocomialis* strain M2 by Clemmer *et al*. [26]. Several studies have reported the involvement of OmpA in biofilm formation [96–98] and OmpA, along with other outer membrane proteins, was observed to accumulate in *A. baumannii* pellicle cells compared to planktonic cells [44]. We found the *ompA* knockout associated with a significant decrease in pellicle biofilm formation in ATCC 17978 but not in 29D2 (Fig. 2). For *A. baumannii* and a number of other pathogens, OmpA has been identified as a virulence factor and its importance in bacterial pathogenicity has been shown recently (reviewed in [99]). For example, the loss of OmpA impaired virulence of *A. baumannii* in *C. elegans* [100] and *Klebsiella pneumoniae* virulence in *G. mellonella* [101]. In our study, the knockout of *ompA* in both tested strains significantly decreased the mutant’s ability to kill *G. mellonella* caterpillars (Fig. 4). We observed 1-2 log scale lower CFU numbers (for both mutant strains) compared to the OD-adjusted suspension which was used to infect the caterpillars. This observation led us to examine the mutant’s cell morphology by microscopy. We found the *ompA::Km* mutants exhibiting filamentous cell phenotypes in contrast to the parental strains (Suppl. Fig. S8). Filamentous cell morphologies are known to provide bacterial survival advantages, e.g. protection against phagocytosis, resistance against antibiotics, and enhanced response to environmental cues like quorum sensing [102]. In other bacteria, the loss of outer membrane proteins, like the Tol-Pal system or OmpA-like proteins, resulted in reduced membrane integrity and alterations in cell division [103, 104]. OmpA is involved in the ability of *A. baumannii* to grow and persist in human serum [11, 105] and in the adherence and invasion of epithelial cells [106]. A resistance phenotype was only observed for the ATCC 17978 *ompA::Km* mutant strain. This mutant showed a 2-fold increase in MIC for imipenem compared to the parental strain (Table 3), correlating with the published finding that the *A. baumannii* OmpA C-terminus is important for resistance to antibiotics including imipenem [107].

### Genes involved in 1,3-diaminopropane biosynthesis

As previously shown, mutations in the genes *dat* and *ddc* resulted in a dramatic reduction in surface-associated motility, but can be restored by supplementation with 120 µM DAP [33]. In the present study, we observed motility deficiency for these genes in 29D2 (Table 2). We also gained new insight into the pleiotropic effects of the *dat::Km* and *ddc::Km* mutants, such as a significant decrease in pellicle biofilm formation. This observation might represent species-specific traits as we see this effect in both tested strains (ATCC 17978 and 29D2), whereas we see contradictory results for the MIC assays (Table 3).

### Genes involved in lipopeptide synthesis/export

The genes *a1s_0113* and *a1s_0116* are involved in the synthesis and export of a lipopeptide and are part of an operon consisting of 8 genes [26, 108]. The knockout of *a1s_0113* (acyl-CoA dehydrogenase) in *A. nosocomialis* clinical isolate M2 resulted in a significant surface motility defect [26], which correlates with our observation in ATCC 17978 (Fig. 1A). We found both mutants to show similar pleiotropic effects in ATCC 17978 (Table 1). Additionally, other genes of this operon have been reported to be necessary for motility (*a1s_0112* and *a1s_0115* [39]), pellicle biofilm formation (*a1s_0112* and *a1s_0115* [39]; *a1s_0114* [108]), and biofilm formation on abiotic surfaces (*a1s_0114* [108, 109]). A pellicle proteome analysis in ATCC 17978 found the proteins A1S_0112-A1S_0118, with the exception of A1S_0114, to accumulate in the pellicle [44]. Since the gene *a1s_0116* encodes an RND superfamily transporter, it may thus play a role in multi-drug resistance. Deletion of *a1s_0116* in ATCC 17978 resulted in significantly increased ampicillin resistance compared to the parental strain whereas no differences were observed for testing with imipenem and tetracycline (Table 3). A transcriptomic study on imipenem-resistant ATCC 17978 cells showed decreased expression of genes from the *a1s_0112-a1s_0119* cluster [110]. Clemmer *et al*. speculated that the lipopeptide synthesized from the *a1s_0112-a1s_0119* operon may act as a surfactant to promote motility, but they could not detect any surfactant activity in *A. nosocomialis* culture supernatants [26]. While we could not show a significant effect of *a1s_0113* or *a1s_0116* inactivation on virulence in *G. mellonella*, significant attenuation was observed in the same model for an *a1s_0114* mutant [108]. No essential role of any of the *a1s_0112*-*a1s_0119* genes in virulence was also found for strain AB5075 [111]. In conclusion, our data confirm findings by other groups [26,39,109,108,44] indicating that genes of the *a1s_0112*-*a1s_0119* operon are essential for surface motility and pellicle biofilm formation in *A. baumannii*.

### Genes involved in DNA modification/repair/uptake

We found 4 genes in our library to be involved in DNA modification, uptake, and recombination. The gene *a1s_0222*, designated as *aamA,* encodes a Type II N6-adenine DNA methyltransferase [112, 113]. Methylation is important for the regulation of various physiological processes [114, 115]. We speculate the phenotype of both *aamA* mutants to represent strain-specific traits (Tables 1 and 2). In bacteria DNA methylation is the most studied epigenetic mechanism and the *E. coli* Dam protein is the most prominent orphan DNA adenine methyltransferase [116]. For other bacteria like *S. enterica*, *Y. enterocolitica,* and *K. pneumoniae* different phenotypes of *dam* mutants and *dam* overexpression were shown to affect motility, virulence, and other traits (reviewed in [117]).

The *A. baumannii* gene *a1s_2334* encodes an S-adenosyl-L-homocysteine hydrolase (*sahH*), which takes part in the recycling of S-adenosyl-L-methionine (SAM). Here we show that inactivation of *sahH* in *A. baumannii* leads to pleiotropic effects such as strong motility deficiency, a significant attenuation in *G. mellonella* caterpillar infection, and increased antibiotic resistance (Table 1). Furthermore, we found the Holliday junction helicase subunit A (*ruvA*/ *a1s_2587*) to be important for *A. baumannii* surface-associated motility, pellicle biofilm formation, and antibiotic resistance in ATCC 17978 (Table 1).

We identified the gene *a1s_2610* in our mutant library screening. Designated as *comEC*, this gene is involved in DNA uptake and incorporation of exogenous DNA into the genome. Phenotypically, a linkage between motility and natural transformation competence was shown in that *A. baumannii* can take up DNA while moving along wet surfaces [16] and its transformability is influenced by motility-determining parameters such as agarose concentration [41]. Genetically this interrelationship was illustrated by abolished twitching motility and natural transformation competence of *comEC* knockout mutants in *A. baumannii* strains 07-095 and 07-102, and a defect in surface-associated motility was ascribed for the ATCC 17978 *comEC::Km* mutant [16]. Deficiency in twitching motility has also been shown for Δ*comEC* in *Thermus thermophilus* [118]. Our results confirmed surface-associated motility deficiency in the 29D2 *comEC::Km* mutant strain (Fig. 2). Deficiency in twitching motility was also shown for Δ*comEC* in *Thermus thermophilus* [118]. A striking attenuation in *G. mellonella* caterpillar infection for the *comEC::Km* mutants in both 17978 and 29D2 was observed (Fig. 4), similar to attenuation of *comEC::Km* mutant derivatives of *A. baumannii* strains DSM 30011, 07-102, and 07-095 [16]. In *Listeria monocytogenes*, *comEC* was demonstrated to be involved in phagosomal escape, intracellular growth, and virulence [119]. However, *com* genes have been reported to be involved in bacterial biofilm formation [120], which we could not confirm for our *comEC::Km* mutant strains (Fig. 2).

### Other genes

The gene *a1s_0065* encodes a UDP-glucose 4-epimerase (*galE*) and is predicted to play a role in capsule and lipopolysaccharide biosynthesis [121]. Capsules are important virulence factors in *A. baumannii* [122]. In this study, a knockout of *galE* resulted in a reduced motility phenotype in ATCC 17978 and 29D2 (Fig. 1). The involvement of lipopolysaccharides in *Acinetobacter* surface motility has recently been shown for the gene *rmlB* which is part of the O-antigen in Gram-negative bacteria [26]. A proteomic study of *A. baumannii* revealed GalE to be only expressed in biofilm growth mode [78]. Additionally, multiple studies have revealed that UDP-glucose 4-epimerases play a role in biofilm formation, including in *Sinorhizobium meliloti* [123], *Vibrio cholerae* [124], *Bacillus subtilis* [125], and *Thermus thermophiles* [126]. The knockout of *galE* resulted in a significant increase in pellicle biomass production for both mutants compared to their parental strains (Fig. 2). Similar observations were also made for a *galE* mutant in *Haemophilus parasuis* [127] and *Porphyromonas gingivalis* [128]. Moreover, other proteins necessary for the catabolism of D-galactose (Leloir pathway), GalM and GalU, were found to be upregulated in *A. baumannii* biofilms [129]. Infection of *G. mellonella* caterpillars with the *galE::Km* mutants resulted in a significant attenuation. The caterpillar survival rate 5 days post infection was 98.9% for the ATCC 17978 *galE::Km* mutant and 95.8% for 29D2 *galE::Km* (Fig. 4). Similarly, a significant decrease in persistence in a mouse pneumonia model of *A. baumannii* was previously reported for the *a1s_0065* mutant [53]. Several other studies demonstrated UDP-glucose 4-epimerases to be involved in virulence/pathogenesis, for example in *Bacillus anthracis* [130], *Streptococcus iniae* [131], and the plant-pathogenic fungus *Leptosphaeria maculans* [132]. In our study, we observed a resistance phenotype for the ATCC 17978 *galE::Km* mutant to ampicillin and tetracycline. By contrast, the 29D2 *galE::Km* mutant showed significant sensitivity to ampicillin compared to the parental strain (Table 3). The *galE* mutant in *Porphyromonas gingivalis* was shown to be significantly more susceptible to benzylpenicillin, oxacillin, cefotaxime, imipenem, and vancomycin compared to the wildtype [128] and involvement of UDP-glucose 4-epimerases in antibiotic resistance/sensitivity was reported for several other bacteria [133–135].

The gene *a1s_0806* encodes for an adenosylmethionine-8-amino-7-oxononanoate aminotransferase (*bioA*), belonging to the acetyl ornithine aminotransferase family, which is part of the pyridoxal phosphate-dependent aspartate aminotransferase superfamily. BioA is part of the biotin biosynthesis pathway and biotin is essential for cell metabolism in prokaryotes and eukaryotes, and only bacteria and plants can synthesize biotin *de novo* [136–138]. Inactivation of *a1s_0806* in ATCC 17978 resulted in the strongest surface motility defect (Fig. 1) and the greatest pellicle biomass production of all tested mutants (Fig. 2). In contrast, the mutant’s ability to kill *G. mellonella* caterpillars was not significant affected (Suppl. Figs. S6 and S7). Other studies in *M. tuberculosis* have demonstrated *bioA* to be essential for establishment of infection and persistence in mice [139].

A knockout of gene *a1s_1055*, encoding a LysM peptidoglycan-binding domain-containing lytic transglycosylase, resulted in a significantly increased pellicle biomass production. Similar to our findings, mutation of lytic transglycosylase (A1S_3027) in *A. nosocomialis* strain M2 was found to exhibit surface motility deficiency [26]. A1S_1055 seems to play a role in *A. baumannii* virulence, since mutants of both parental backgrounds led to attenuation in the *G. mellonella* infection assay (Fig. 4). The gene *a1s_2761* encodes for a 2-methylaconitate cis-trans isomerase (PrpF), involved in the 2-methylcitric acid cycle and propionate catabolism. The inactivation of *prpF* in 17978 resulted in one of the most reduced pellicle formation phenotypes of all tested mutant strains (Fig. 2), which correlates with the previous finding that PrpF accumulated in mature 4-days pellicles in 17978 [44]. The mutant’s virulence capacity was not significantly affected (Suppl. Fig. S6). In contrast, the 29D2 *prpF::Km* mutant strain displayed a significant attenuation in the *G. mellonella* infection model (Fig. 4) but did not affect pellicle biofilm formation (Fig. 2).

The *A. baumannii* gene *a1s_3026* is predicted to encode a secreted ribonuclease T2 family protein (RNase T2-family). In the *A. baumannii* strain 98-37-09 a deficiency in surface motility for the *a1s_3026* knockout was previously reported [34]. Additionally, Jacobs *et al*. indicated that the *a1s_3026* mutant showed reduced colonization on abiotic surfaces like glass, polystyrene, and stainless steel, and that the *a1s_3026* knockout was shown to be associated with decreased expression of genes involved in motility and biofilm formation [34]. Despite the deficiency in surface-associated motility we observed a significant decrease in pellicle biofilm formation (Tables 1 and 2). Interestingly, A1S_3026 was shown to be involved in *A. baumannii* colistin resistance [140] and both *3026::Km* mutant strains exhibited elevated resistance values for ampicillin and imipenem (Table 3).

The *A. baumannii* gene *a1s_3129* encodes for a succinylarginine dihydrolase (*astB*) and is involved in the arginine succinyltransferase (AST) pathway [141]. In a mouse pneumonia model of *A. baumannii*, the *astB* insertion caused a significant decrease in persistence [53]. However, we did not observe a significant attenuation in the *G. mellonella* infection model (Table 1). Furthermore, we observed a significant reduction of pellicle formation in the *astB::Km* mutant (Fig. 2A). This is in line with an accumulation of AstB in ATCC 17978 pellicle cells described previously [44].

### *G. mellonella* caterpillar infection

The *G. mellonella* caterpillar, which is an established insect model system for bacterial infections [142], was used to study virulence traits of the motility-deficient mutants. A study on virulence and resistance to antibiotic and environmental stress analyzed 250,000 *A. baumannii* AB5075 transposon mutants for growth within *G. mellonella* larvae, and TnSeq experiments identified 300 genes essential for growth [111]. When comparing with these results, we could not identify concordant genes in our library, but we found that main categories of genes do match. For example we found *galE* to be essential and in AB5075 numerous genes involved in structure and function of the cell envelopment were found to be required for growth in *G. mellonella* [111]. Conversely, for example, the *gidA∷Km* mutant was not attenuated in *G. mellonella* infection in our study (Suppl. Fig. S6 and Fig. S7), but was stated to be essential for growth of AB5075 in *G. mellonella* [111]. It is known that AB5075 is more virulent than ATCC 17978 [143, 111], therefore, comparative studies are needed to unravel strain-specific and species-specific traits.

### Limitations

While our study highlights the need for comparative studies of specific mutant phenotypes in different strains to distinguish strain-specific from species-specific traits, it is clear that the two strains studied in detail here do not provide a sufficient basis to deduce such insight. Such comparative studies in combination with genome-based analyses may pave the way for the identification of species-specific traits and, ultimately, novel target sites.

The use of marker-based mutagenesis and naturally competent strains to efficiently generate sets of mutants in different strains has its shortcomings as recombination events are not necessarily limited to the site of the marker gene. Apart from homology-based recombination events, transfer of mobile genetic elements and even illegitimate recombination events may occur [144, 145] and could corrupt the mutants’ phenotypes. A few of the mutations described in this study have been partially characterized previously using additional strains (*ddc*, *dat*, *comEC,* and *aamA* for example [16,33,112]). However, repetitive construction of the same mutants did not lead to significant phenotype variation arguing against a high frequency of such corrupting side-effects. The ability to rapidly reconstruct mutants is an advantage of this method which can allow for confirmation of mutant phenotypes.

Complementation experiments and site-specific deletion mutagenesis would exclude polar effects of the transposon insertions and help to verify the contribution of each gene. In support of the specificity of our findings, we found many groups of related mutants (e.g. purine and pyrimidine biosynthesis) and identified multiple linkages to motility mutants described in other organisms.

We could not achieve a saturated mutant library which indicates that surface-associated motility is probably under control of additional genes yet to be discovered. Our attempts were limited by the poor transformability of ATCC 17978 with the transposome complex, and by the transposition not being completely unbiased so that we obtained several insertion events repeatedly and independently.

## Conclusion

In this study we made use of a previously generated *A. baumannii* ATCC 17978 EZ-Tn5™ <KAN-2> transposon mutant library [33] to screen for surface-associated motility-deficient mutants. We identified 30 genes involved in surface motility. All tested mutants originally identified as motility-deficient in strain ATCC 17978 also displayed a motility-deficient phenotype in the *A. baumannii* white stork isolate 29D2. Some of these genes have already been linked to motility in *A. baumannii* (e.g. *comEC*, *a1s_0113,* and *a1s_0116*) or other bacteria (e.g. *carB* and *gidA*), but some of our findings represent new insights into requirements for surface-associated motility. Furthermore, we analyzed these mutants with respect to bacterial growth, pellicle biofilm formation, virulence in *G. mellonella* infection, and antibiotic resistance and used the naturally competent strain 29D2 to indicate whether the mutations showed strain-specific or species-specific traits. In summary, we can state that mutations in genes involved in purine/pyrimidine/folate biosynthesis are essential for all tested categories. Mutants that targeted RNA modification/regulation seem to mainly play a role in motility and pellicle formation. The discovery of novel genes required for surface-associated motility in *A. baumannii* demonstrates that more work is required to further define its genetic basis.

## Conflict of Interest

The authors declare that they have no conflict of interest.

