## Supplementary material for "Novel genes required for surface-associated motility in ***Acinetobacter baumannii***"

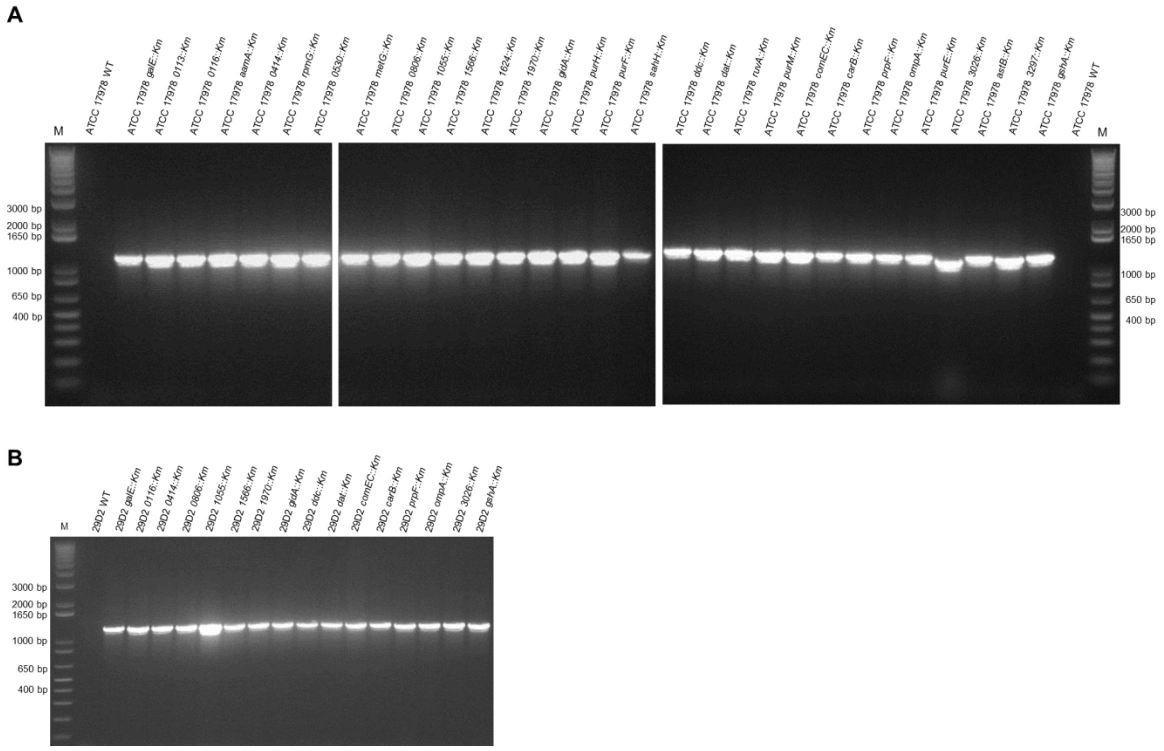


**Fig. S1. PCR confirmation of EZ-Tn5™ <KAN-2> transposon insertion in ATCC 17978 (A) and 29D2 (B) mutants.** PCR using the kanamycin cassette primers of the EZ-Tn5™ <KAN-2> insertion kit (Epicentre Biotechnologies). Insertion of the EZ-Tn5™ <KAN-2> transposon results in a 1221 bp PCR product. Both wildtype strains, ATCC 17978 WT (A) and 29D2 WT (B), are used as a negative control.

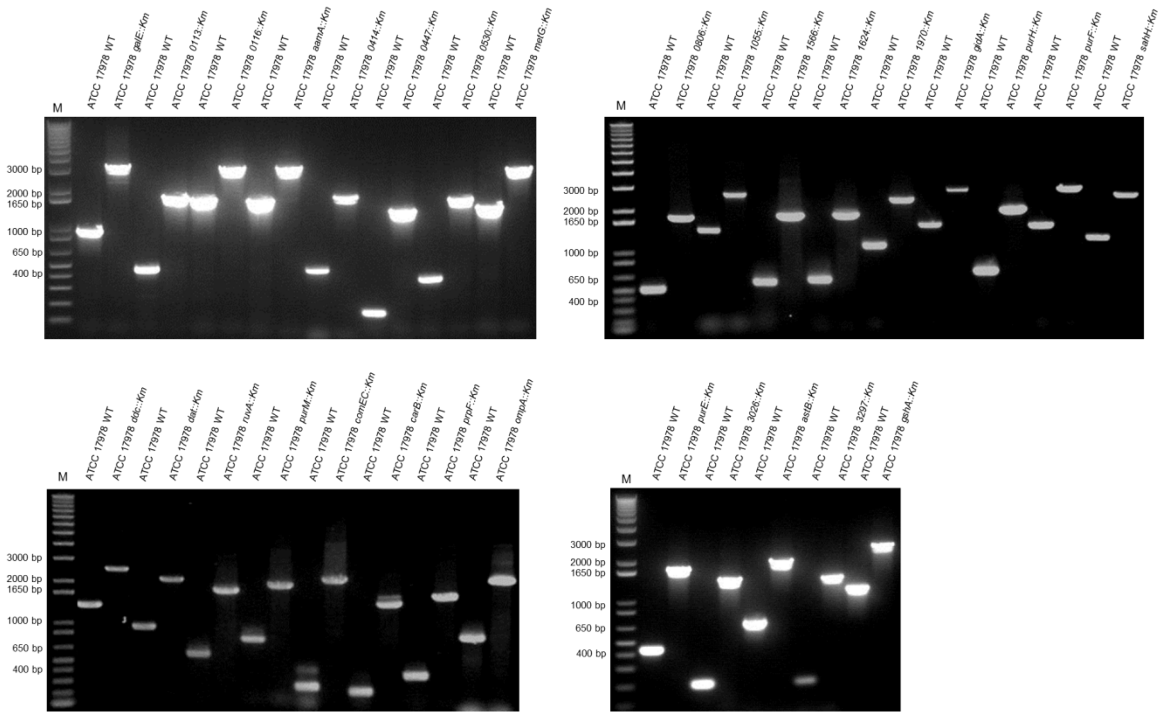


**Fig. S2. PCR confirmation of specific gene inactivation in strain ATCC 17978.** For each mutant, specific gene target site primers are used as described in Materials & Methods and Table S2. The expected size of PCR products for each mutant is given in Table S2.

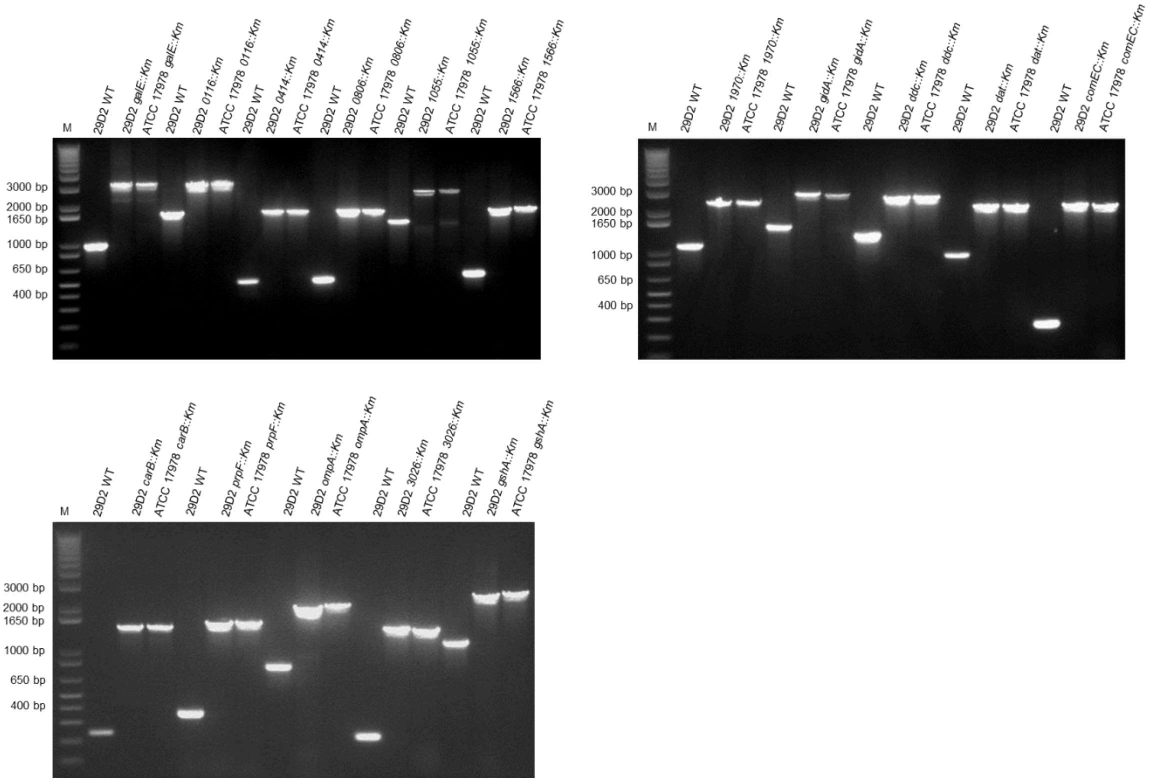


**Fig. S3. PCR confirmation of specific gene inactivation in strain 29D2.** Appropriate gene target site primers are used as described in Materials & Methods and Table S2. Defined length of PCR products for each mutant is given in Table S2.

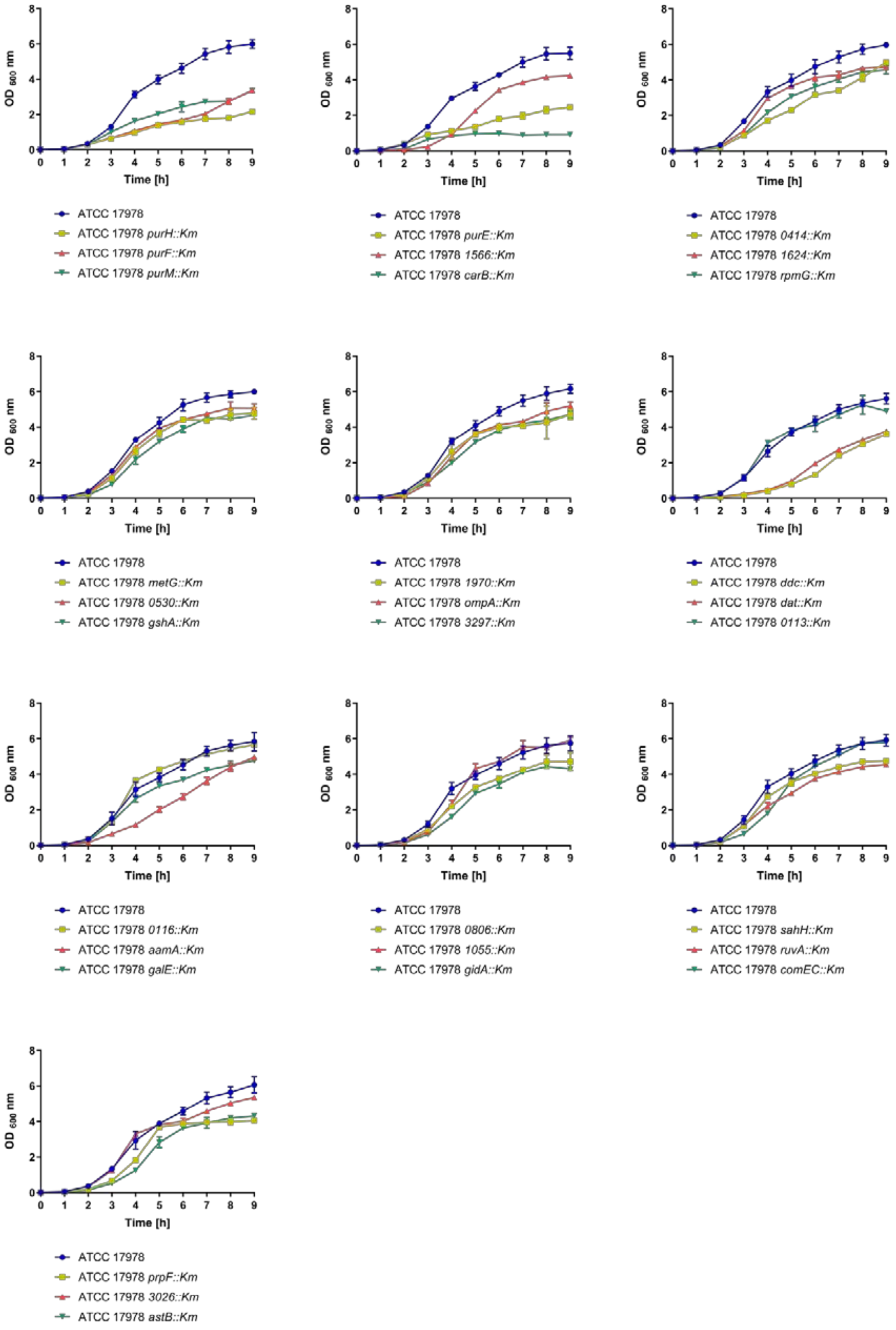


**Fig. S4. Growth curves of ATCC 17978 wildtype and mutant strains.** OD-adjusted bacterial cultures were incubated for 9 hours in baffled flasks at 37°C under constant shaking. Every hour cultures were measured at an OD of 600 nm. For each strain data obtained from three independent cultures grown on the same day were averaged and represented by the mean ± SD. The ATCC 17978 wildtype is indicated by a blue line.

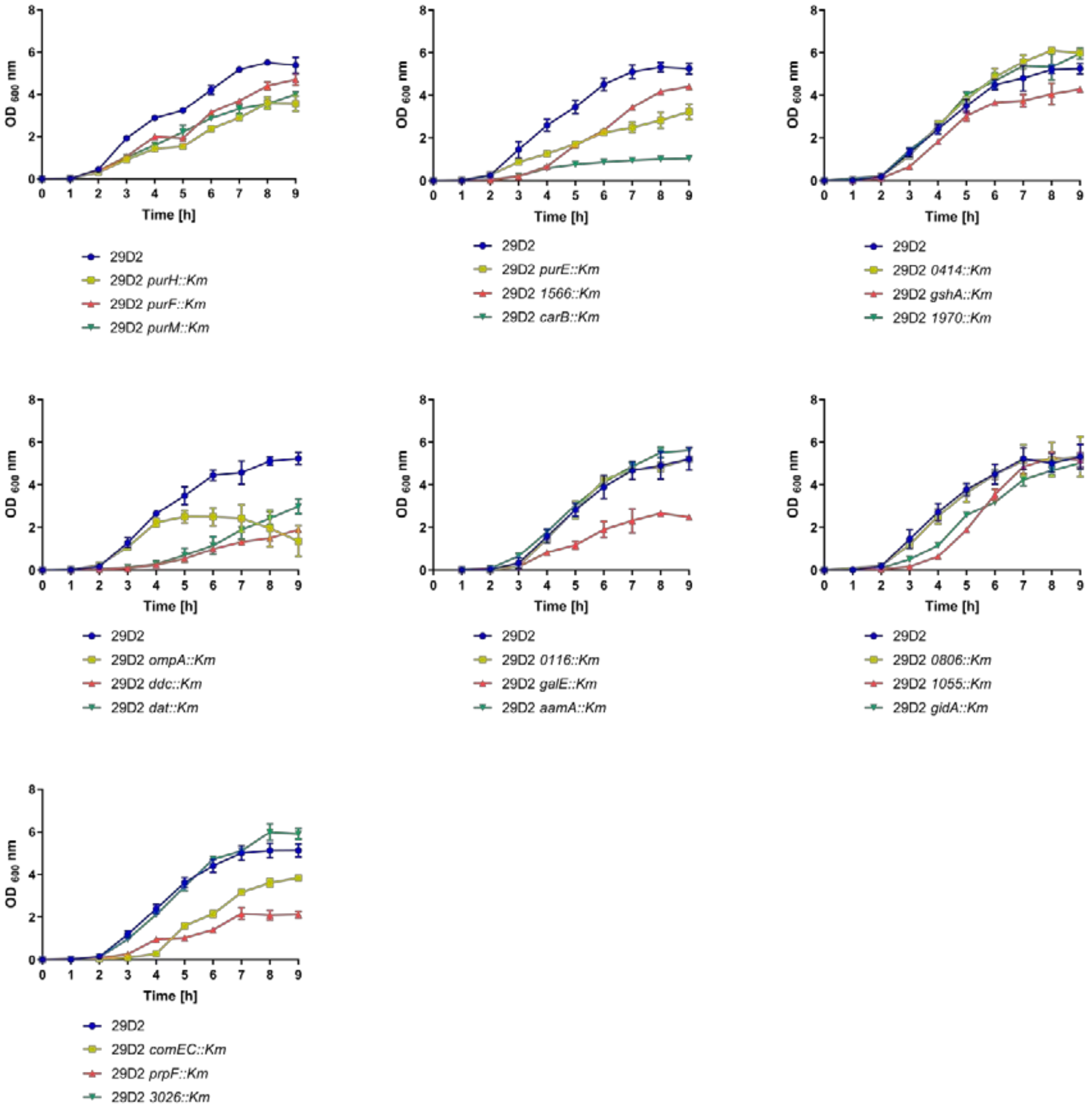


**Fig. S5. Growth curves of 29D2 wildtype strain and 29D2 mutants.** OD adjusted overnight cultures were incubated for 9 hours in baffled flasks at 37°C under constant shaking. Every hour cultures were measured at an OD of 600 nm. For each strain data obtained from three independent cultures grown on the same day were averaged and represented by the mean ± SD. The 29D2 wildtype is indicated by a blue line.

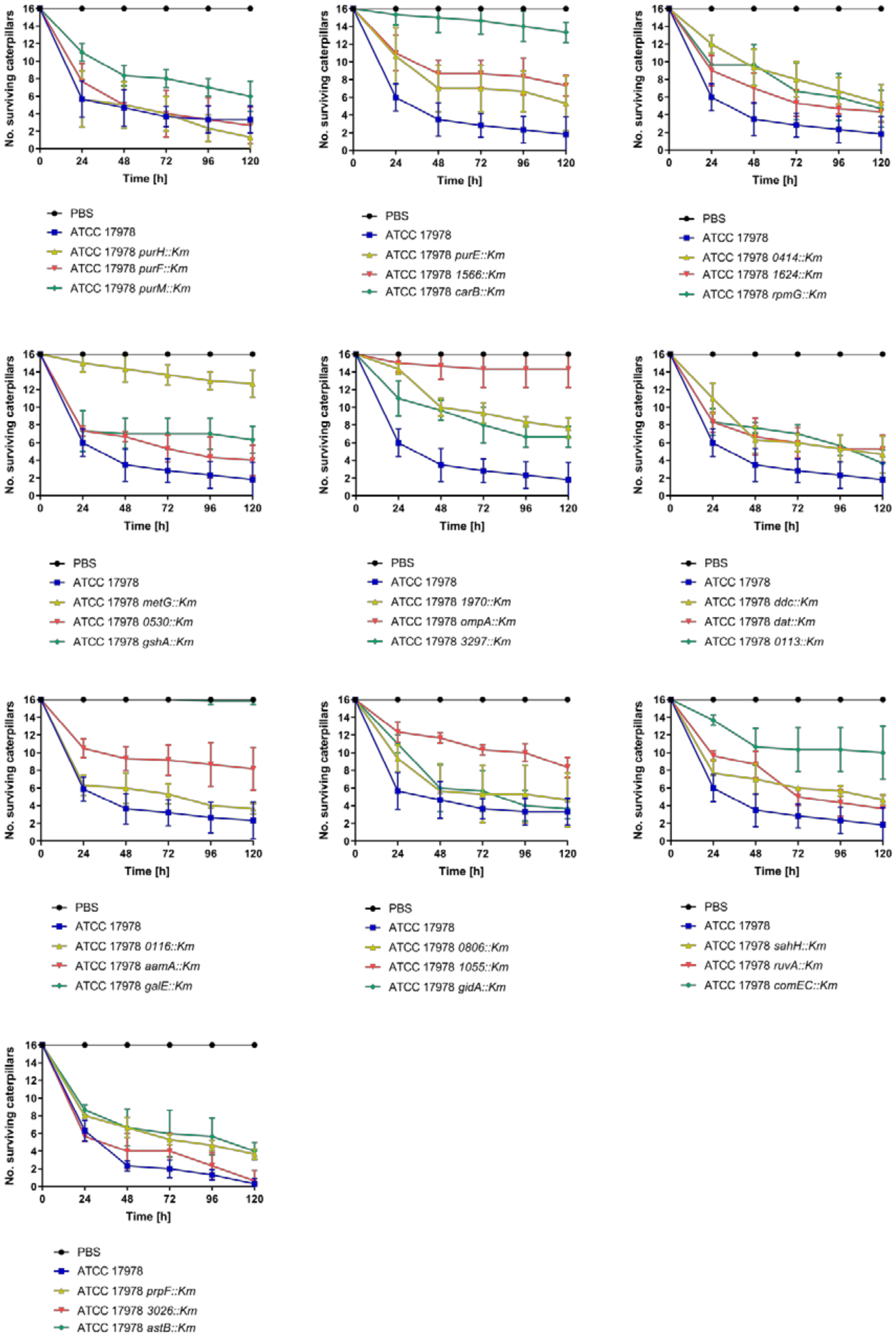


**Fig. S6. *Galleria mellonella* caterpillars infected with ATCC 17978 wildtype and mutant strains.** Caterpillars were infected with 3 x $10^5$ CFU of either ATCC 17978

wildtype (blue line) or mutant strains. Sterile PBS was used as a control (black line). Three independent experiments were performed with groups of 16 caterpillars for every bacteria strain and control. Data obtained from three independent experiments were averaged and represented by the mean ± SD. Significant attenuation in caterpillar infection is observed for mutants *carB::Km*, *ompA::Km*, *metG::Km* and *galE::Km* after 5 days post infection.

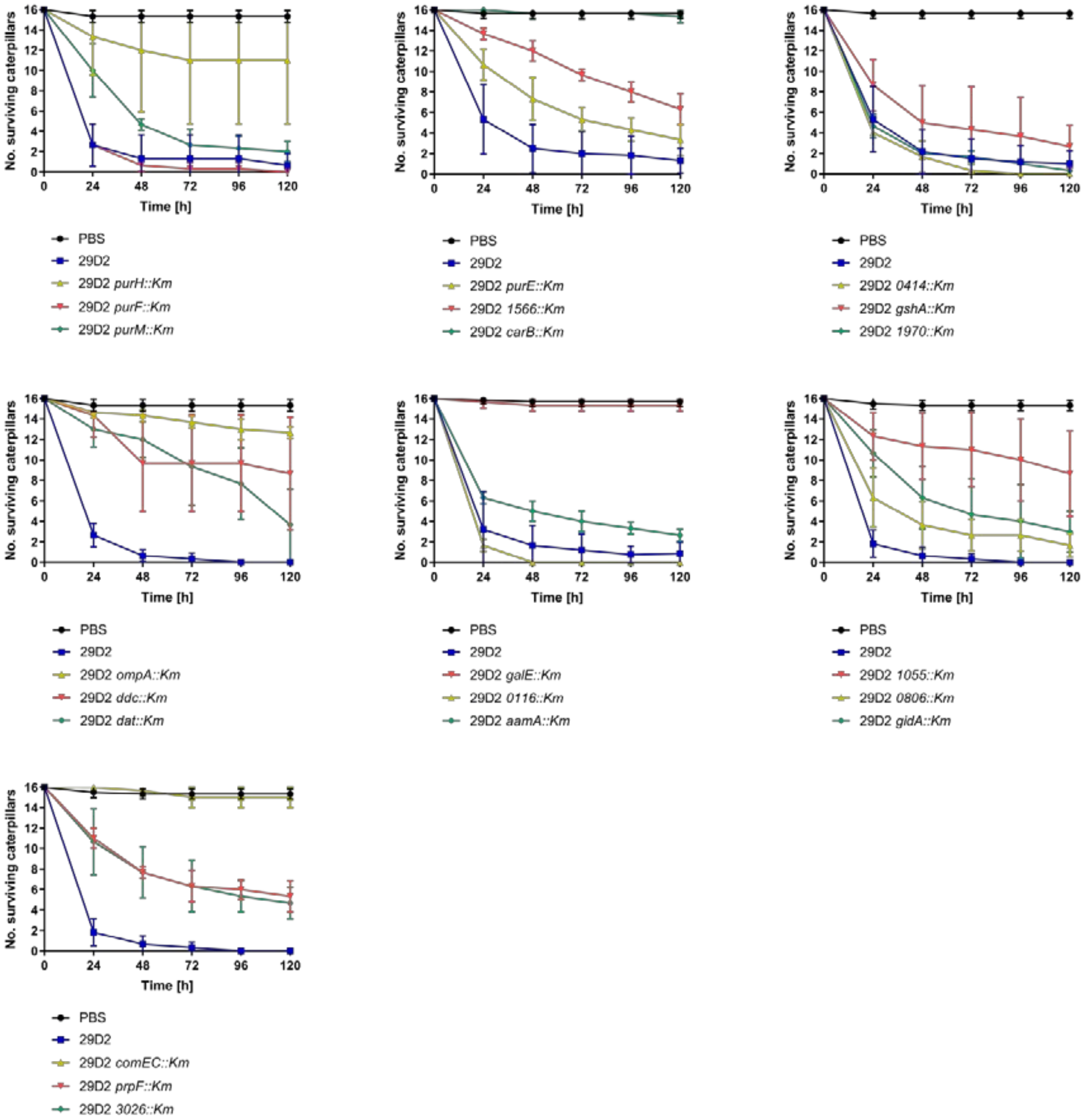


**Fig. S7. Infection of *Galleria mellonella* caterpillars with 29D2 wildtype and 29D2 mutant strains.** *G. mellonella* caterpillars were infected with 3 x $10^5$ CFU of 29D2 wildtype (blue line) or mutant strains. As a control sterile PBS was used (black line). Three independent experiments were performed with groups of 16 caterpillars for every bacterial strain and control. Data obtained from three independent experiments were averaged and represented by the mean ± SD. Significant attenuation after 5 days p.i. is observed for mutants *carB::Km*, *ompA::Km*, *galE::Km* and *comEC::Km.*

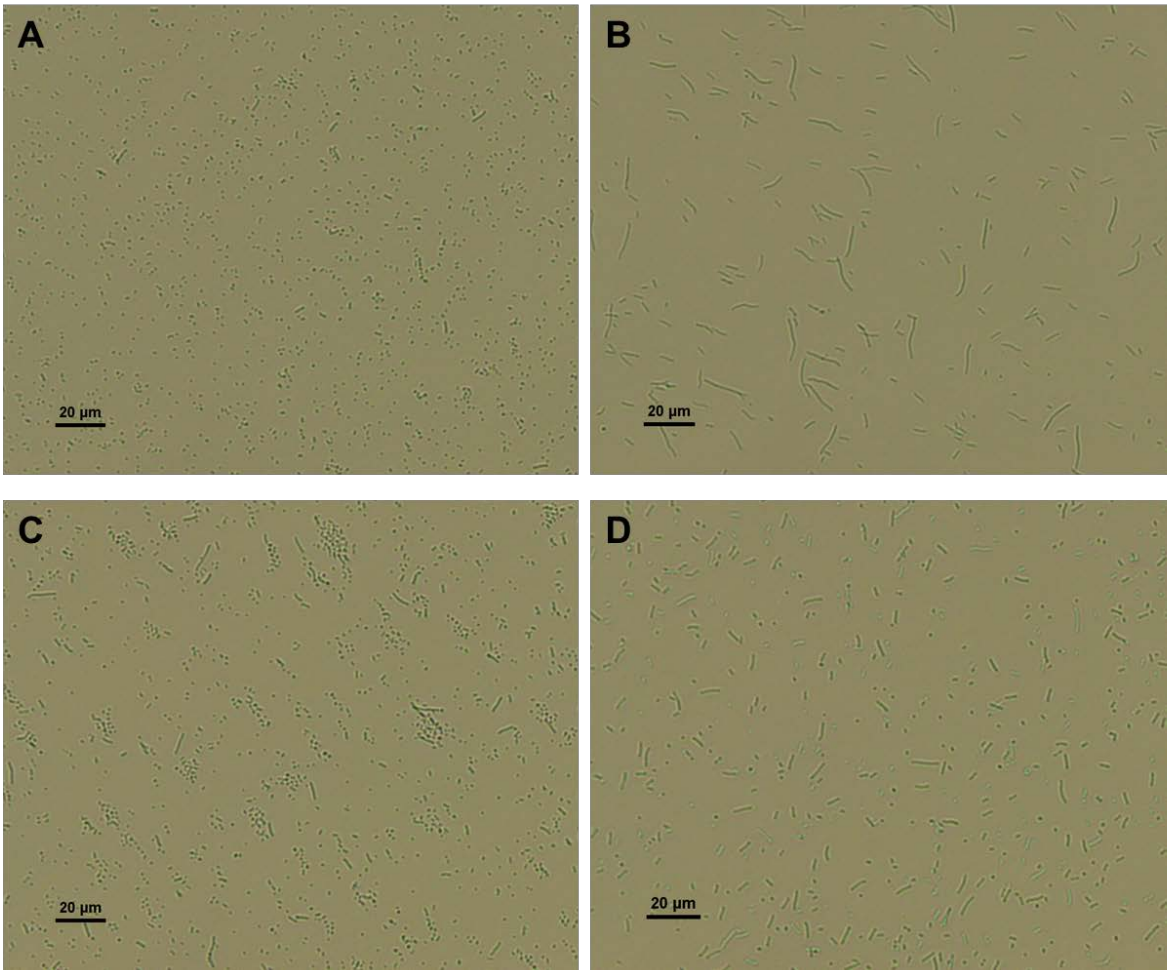


**Fig. S8. Bright field microscopy of *ompA::Km* mutants (B,D) and parental strains (A,C).** Bacterial cells were grown for 16 hours at 37°C under constant shaking. One microliter of each bacterial strain was pipetted on a glass slide and analyzed under the bright field microscope (200-fold magnification). The ATCC 17978 wildtype strain **(A)** and 29D2 wildtype strain **(C)** display round to rod-shaped bacterial cells. Both, the ATTC 17978 *ompA::Km* mutant **(B)** and the 29D2 *ompA::Km* mutant **(D)** display an elongated or chain structure phenotype.

### Table S1. Bacterial strains used in this work

| **Bacteria strain** | **Genotype** | **Source** |
|---|---|---|
| *A. baumannii* ATCC 17978 | wildtype strain | American Type Culture Collection (ATCC) |
| ATCC 17978 *0065::Km* | A1S_0065::EZ-Tn5™ <KAN-2> (*galE*) | This work |
| ATCC 17978 *0113::Km* | A1S_0113::EZ-Tn5™ <KAN-2> | This work |
| ATCC 17978 *0116::Km* | A1S_0116::EZ-Tn5™ <KAN-2> | This work |
| ATCC 17978 *aamA::Km* | A1S_0222::EZ-Tn5™ <KAN-2> (*aamA*) | [110] |
| ATCC 17978 *0414::Km* | A1S_0414::EZ-Tn5™ <KAN-2> | This work |
| ATCC 17978 *0447::Km* | A1S_0447::EZ-Tn5™ <KAN-2> (*rpmG*) | This work |
| ATCC 17978 *0530::Km* | A1S_0530::EZ-Tn5™ <KAN-2> | This work |
| ATCC 17978 *0778::Km* | A1S_0778::EZ-Tn5™ <KAN-2> (*metG*) | This work |
| ATCC 17978 *0806::Km* | A1S_0806::EZ-Tn5™ <KAN-2> | This work |
| ATCC 17978 *1055::Km* | A1S_1055::EZ-Tn5™ <KAN-2> | This work |
| ATCC 17978 *1566::Km* | A1S_1566::EZ-Tn5™ <KAN-2> | This work |
| ATCC 17978 *1624::Km* | A1S_1624::EZ-Tn5™ <KAN-2> | This work |
| ATCC 17978 *1970::Km* | A1S_1970::EZ-Tn5™ <KAN-2> | This work |
| ATCC 17978 *gidA::Km* | A1S_2182::EZ-Tn5™ <KAN-2> (*gidA*) | This work |
| ATCC 17978 *purH::Km* | A1S_2187::EZ-Tn5™ <KAN-2> (*purH*) | This work |
| ATCC 17978 *purF::Km* | A1S_2251::EZ-Tn5™ <KAN-2> (*purF*) | This work |
| ATCC 17978 *sahH::Km* | A1S_2334::EZ-Tn5™ <KAN-2> (*sahH*) | This work |
| ATCC 17978 *ddc::Km* | A1S_2453::EZ-Tn5™ <KAN-2> (*ddc*) | [33] |
| ATCC 17978 *dat::Km* | A1S_2454::EZ-Tn5™ <KAN-2> (*dat*) | [33] |
| ATCC 17978 *2587::Km* | A1S_2587::EZ-Tn5™ <KAN-2> (*ruvA*) | This work |
| ATCC 17978 *purM::Km* | A1S_2605::EZ-Tn5™ <KAN-2> (*purM*) | This work |
| ATCC 17978 *comEC::Km* | A1S_2610::EZ-Tn5™ <KAN-2> (*comEC*) | [16] |
| ATCC 17978 *2687::Km* | A1S_2687::EZ-Tn5™ <KAN-2> (*carB*) | This work |
| ATCC 17978 *prpF::Km* | A1S_2761::EZ-Tn5™ <KAN-2> (*prpF*) | This work |

| | | |
|---|---|---|
| ATCC 17978 *ompA::Km* | A1S_2840::EZ-Tn5™ <KAN-2> (*ompA*) | This work |
| ATCC 17978 *purE::Km* | A1S_2964::EZ-Tn5™ <KAN-2> (*purE*) | This work |
| ATCC 17978 *3026::Km* | A1S_3026::EZ-Tn5™ <KAN-2> | This work |
| ATCC 17978 *astB::Km* | A1S_3129::EZ-Tn5™ <KAN-2> (*astB*) | This work |
| ATCC 17978 *3297::Km* | A1S_3297::EZ-Tn5™ <KAN-2> | This work |
| ATCC 17978 *3366::Km* | A1S_3366::EZ-Tn5™ <KAN-2> (*gshA*) | This work |
| *A. baumannii* 29D2 | wildtype strain | White stork isolate [39] |
| 29D2 *aamA::Km* | A1S_0222::EZ-Tn5™ <KAN-2> (*aamA*) | This work |
| 29D2 *0065::Km* | A1S_0065::EZ-Tn5™ <KAN-2> (*galE*) | This work |
| 29D2 *0116::Km* | A1S_0116::EZ-Tn5™ <KAN-2> | This work |
| 29D2 *0414::Km* | A1S_0414::EZ-Tn5™ <KAN-2> | This work |
| 29D2 *0806::Km* | A1S_0806::EZ-Tn5™ <KAN-2> | This work |
| 29D2 *1055::Km* | A1S_1055::EZ-Tn5™ <KAN-2> | This work |
| 29D2 *1566::Km* | A1S_1566::EZ-Tn5™ <KAN-2> | This work |
| 29D2 *1970::Km* | A1S_1970::EZ-Tn5™ <KAN-2> | This work |
| 29D2 *gidA::Km* | A1S_2182::EZ-Tn5™ <KAN-2> (*gidA*) | This work |
| 29D2 *ddc::Km* | A1S_2453::EZ-Tn5™ <KAN-2> (*ddc*) | This work |
| 29D2 *dat::Km* | A1S_2454::EZ-Tn5™ <KAN-2> (*dat*) | This work |
| 29D2 *comEC::Km* | A1S_2610::EZ-Tn5™ <KAN-2> (*comEC*) | This work |
| 29D2 *2687::Km* | A1S_2687::EZ-Tn5™ <KAN-2> (*carB*) | This work |
| 29D2 *prpF::Km* | A1S_2761::EZ-Tn5™ <KAN-2> (*prpF*) | This work |
| 29D2 *ompA::Km* | A1S_2840::EZ-Tn5™ <KAN-2> (*ompA*) | This work |
| 29D2 *3026::Km* | A1S_3026::EZ-Tn5™ <KAN-2> | This work |
| 29D2 *3366::Km* | A1S_3366::EZ-Tn5™ <KAN-2> (*gshA*) | This work |
| 29D2 *purH::Km* | A1S_2187::EZ-Tn5™ <KAN-2> (*purH*) | This work |
| 29D2 *purF::Km* | A1S_2251::EZ-Tn5™ <KAN-2> (*purF*) | This work |
| 29D2 *purM::Km* | A1S_2605::EZ-Tn5™ <KAN-2> (*purM*) | This work |
| 29D2 *purE::Km* | A1S_2964::EZ-Tn5™ <KAN-2> (*purE*) | This work |

### Table S2. Oligonucleotides used for determination of gene target sites

| Primer name | Forward primer (3‘ → 5‘) | Reverse primer (5‘ → 3‘) | PCR product in wildtype strain [bp][a] | Locus tag in ATCC 17978 |
|---|---|---|---|---|
| 0065 for/rev | GCAGGTTATATTGGTTCACACAC | CCATTAGGATTCTGCTTTTGCC | 980 | A1S_0065 |
| Acyl-CoA-DH for/rev | ATGTCACTTACGTCCAATTTGCG | CGATTGATCGACCATTTCTCACC | 488 | A1S_0113 |
| 0116 for/rev | TGACCATAGAACATGCTTTCCTC | CTTTGCGTTTGACCTTCACAACC | 1668 | A1S_0116 |
| BamHI-0222 for/rev | GGATCCGGATGAAATGATCAGTTATGTGGC | GGATCCGTGAGACAGATCCCGTTAGTTGC | 1649 | A1S_0222 |
| 313/0414 for/rev | TAATTGTTGCACTGGCAACTGTG | TGATCGACTCAGAAGGTTTCCG | 479 | A1S_0414 |
| 0447 for/rev | GATTTTAGCTTCTTTGAAAATCACG | ATGCGTGATAAGATTCGCCTCG | 150 | A1S_0447 |
| 0530 for/rev | TTTTGGTTTGCTTTTAACGAGAGG | GGAACGCTGGTTAGAGTTTATGG | 409 | A1S_0530 |
| 0778 for/rev | CTTTATCGCCAGGTTTTGCACC | CCTCTTTCAATTGCTAACAAATTGG | 1432 | A1S_0778 |
| AT_0806 for/rev | GCAACGATTGAACTTGATGATGG | CTGAATAGCTTGCTCTAGCTCG | 498 | A1S_0806 |
| MTG_1055 for/rev | ATCCACGTATTGAAGCACAGCG | TACTGTATAAGTACTACGCTTACC | 1427 | A1S_1055 |
| 1566 for/rev | GTTTGAAAATGCACACGTTGTACG | CTTCTACTTGTAAGTTAACATGCG | 569 | A1S_1566 |
| 1624 for/rev | TTGCATCCACAACTTGCTCAAG | CGGTGAGGTAAAGGCATAGATC | 592 | A1S_1624 |
| 1970 for/rev | ATCACTTTTTACACCAAGTACTCC | TAATGAAATTGCAGTTGAAGCATTC | 1103 | A1S_1970 |
| GidA_2182 for/rev | GGACGTTTAAGCAAATCAATCGC | CTATCCTAAAGTTTATGATGTTATCG | 1542 | A1S_2182 |
| 2187 for/rev | GCCGTGTTAAAACACTACATCC | TCTAAAGAAACGGCAACACCAC | 685 | A1S_2187 |

| | | | | |
|---|---|---|---|---|
| APT_2251 for/rev | CGGTCAAATCTACTGATGCTGC | AGTGAACCAAATGTTGTTTGATGC | 1490 | A1S_2251 |
| 2334 for/rev | AGATTACAAAGTTGCTGACATCTC | ACTTCTACGCGAATTTTTGCAGC | 1212 | A1S_2334 |
| 2453 for/rev | CGTAGTTACCAGAAGCAATCGC | CGTTCAGCAATTTCTTTCCAACC | 1363 | A1S_2453 |
| AT_2454 for/rev | CACGGAAAGTACCAGTATGACC | TGAGCGTTACTTCTGTCAACCC | 957 | A1S_2454 |
| 2587 for/rev | ATGTTTAATTGGCGAAGTGTTTGC | TACTTCATCATTGATTTAAGTGCAGC | 591 | A1S_2587 |
| 2605 for/rev | CCGGTTTAAGCTACAAAGATGCG | AGCTTCATTAATTACAGCTTGAGC | 800 | A1S_2605 |
| 134/2610 for2 / 2610-comb rev | CAATTAGCAACAGTAGACTTGCG | CCCACATGTGCTCATTTTTGCC | 278 | A1S_2610 |
| 172/2687 for/rev | GTTCGTGACAAAAACGACAACTG | TTCATCTCGATCACAACCATACG | 242 | A1S_2687 |
| 120/2761 for/rev | CCAGATTCGAACAGTACAGAGGC | CAGCAACTTACATGCGTGGTGG | 383 | A1S_2761 |
| OmpA for/rev | TGAGCTGCTGCAGGAGCTGC | AAGTTAAAGGCGACGTAGACGG | 826 | A1S_2840 |
| 188/2964 for/rev | CCTGCCCCGGAATATTGTTGC | ATGGGTTCCCAGTCCGATTGG | 463 | A1S_2964 |
| Klon1-RNaseT2 for/rev | GCTTGGGCAATTAGAGACAACG | AAAATACCTGCTAATCTAACCAGC | 222 | A1S_3026 |
| 25/3129 for/rev | GACTCTTCAGGCACAACAATGG | CGGAGATGACATTAGTGTTGTGG | 756 | A1S_3129 |
| 130/3297 for/rev | TTTACTTAACTCCTAAGTTAAGTGTTGG | AGCCGATAGTTTGTGTATCGTAGC | 204 | A1S_3297 |
| 354/3366 for/rev | TTAGAAATATGCTGACGTGCCC | CGTGGAATAGAACGTGAAAGCC | 1179 | A1S_3366 |

[a] Based on the insertion of the EZ-Tn5™ <KAN-2> transposon (1221 bp), PCR products in mutant strains are 1221 bp longer than PCR products from wildtype strains

**Table S3. Mean and standard deviation (SD) of ATCC 17978 wildtype/mutants and 29D2 wildtype/mutants surface motility spreading zones**

| Locus tag | Gene name | Mean ± SD diameter of surface motility spreading zone [mm][a] | |
|---|---|---|---|
| | | ATCC 17978 | 29D2 |
| **Wildtype** | | 77.5 ± 10.6 | 30 ± 10 |
| **A1S_2187** | *purH* | 10.25 ± 3.5 | 5 ± 1.15 |
| **A1S_2251** | *purF* | 6 ± 0 | 3.75 ± 0.95 |
| **A1S_2605** | *purM* | 3.62 ± 1.10 | 16 ± 1.63 |
| **A1S_2964** | *purE* | 3.5 ± 1.29 | 6.25 ± 1.25 |
| **A1S_1566** | | 3.75 ± 0.5 | 10 ± 2.94 |
| **A1S_2687** | *carB* | 4.5 ± 2.38 | 7.75 ± 4.19 |
| **A1S_0414** | | 4.37 ± 1.10 | 12.25 ± 2.62 |
| **A1S_1624** | | 7.25 ± 0.95 | - |
| **A1S_0447** | *rpmG* | 3.75 ± 1.70 | - |
| **A1S_0778** | *metG* | 2.62 ± 0.47 | - |
| **A1S_0530** | | 5 ± 1.82 | - |
| **A1S_3366** | *gshA* | 4.875 ± 1.54 | 6.5 ± 1.29 |
| **A1S_1970** | | 8.75 ± 4.27 | 8.75 ± 3.40 |
| **A1S_2840** | *ompA* | 6.5 ± 4.04 | 10.5 ± 1 |

| | | | |
|---|---|---|---|
| **A1S_3297** | | 11 ± 2.16 | - |
| **A1S_2453** | *ddc* | 4 ± 1.41 | 4.25 ± 1.25 |
| **A1S_2454** | *dat* | 3 ± 1.82 | 7.75 ± 2.06 |
| **A1S_0113** | | 4.75 ± 0.95 | - |
| **A1S_0116** | | 5 ± 1.82 | 10 ± 4.89 |
| **A1S_0222** | *aamA* | 4.75 ± 2.75 | 7.75 ± 3.30 |
| **A1S_0065** | *galE* | 4.5 ± 1.29 | 5.5 ± 1.91 |
| **A1S_0806** | | 1 ± 0.81 | 7 ± 1.15 |
| **A1S_1055** | | 5 ± 0.81 | 8.25 ± 2.06 |
| **A1S_2182** | *gidA* | 4.25 ± 0.5 | 7.75 ± 0.5 |
| **A1S_2334** | *sahH* | 3.75 ± 1.70 | - |
| **A1S_2587** | *ruvA* | 5 ± 1.41 | - |
| **A1S_2610** | *comEC* | 5.5 ± 2.38 | 6.25 ± 1.70 |
| **A1S_2761** | *prpF* | 4 ± 1.41 | 9.5 ± 0.57 |
| **A1S_3026** | | 4.87 ± 0.25 | 9 ± 2.58 |
| **A1S_3129** | *astB* | 3 ± 0.40 | - |

[a] For each strain three independent experiments were performed and represented by the mean ± SD

**Table S4. Mean and standard deviation (SD) of ATCC 17978 wildtype/mutants and 29D2 wildtype/mutants pellicle biofilm measurements**

| Locus tag | Gene name | Mean ± SD [OD $_{550\ nm}$][a] | |
|---|---|---|---|
| | | ATCC 17978 | 29D2 |
| **Wildtype** | | 8.602 ± 1.519 | 5.560 ± 1.330 |
| **A1S_2187** | *purH* | 11.403 ± 1.540 | 7.266 ± 0.716 |
| **A1S_2251** | *purF* | 6.786 ± 1.065 | 5.98 ± 0.128 |
| **A1S_2605** | *purM* | 10.663 ± 0.831 | 4.793 ± 0.902 |
| **A1S_2964** | *purE* | 11.883 ± 0.387 | 4.71 ± 0.918 |
| **A1S_1566** | | 1.806 ± 0.969 | 7.303 ± 3.210 |
| **A1S_2687** | *carB* | 0.570 ± 0.195 | 3.166 ± 0.707 |
| **A1S_0414** | | 0.343 ± 0.211 | 5.156 ± 1.207 |
| **A1S_1624** | | 7.95 ± 1.063 | - |
| **A1S_0447** | *rpmG* | 9.943 ± 2.089 | - |
| **A1S_0778** | *metG* | 2.259 ± 0.397 | - |
| **A1S_0530** | | 13.1 ± 1.375 | - |
| **A1S_3366** | *gshA* | 9.573 ± 1.160 | 5.976 ± 0.206 |
| **A1S_1970** | | 5.633 ± 1.896 | 3.136 ± 0.770 |
| **A1S_2840** | *ompA* | 1.064 ± 0.609 | 3.93 ± 2.170 |
| **A1S_3297** | | 11.093 ± 1.165 | - |

| | | | |
|---|---|---|---|
| **A1S_2453** | *ddc* | 3.213 ± 1.868 | 2.606 ± 0.669 |
| **A1S_2454** | *dat* | 2.308 ± 1.619 | 3.203 ± 0.090 |
| **A1S_0113** | | 5.726 ± 0.781 | - |
| **A1S_0116** | | 5.283 ± 1.504 | 4.196 ± 0.342 |
| **A1S_0222** | *aamA* | 4.123 ± 2.237 | 6.748 ± 1.626 |
| **A1S_0065** | *galE* | 14.943 ± 2.844 | 7.723 ± 1.337 |
| **A1S_0806** | | 15.88 ± 2.104 | 5.963 ± 0.601 |
| **A1S_1055** | | 11.4 ± 1.018 | 7.126 ± 0.977 |
| **A1S_2182** | *gidA* | 2.556 ± 0.295 | 3.883 ± 0.261 |
| **A1S_2334** | *sahH* | 9.95 ± 0.645 | - |
| **A1S_2587** | *ruvA* | 4.4 ± 1.368 | - |
| **A1S_2610** | *comEC* | 8.813 ± 3.092 | 4.486 ± 0.680 |
| **A1S_2761** | *prpF* | 0.687 ± 0.405 | 6.21 ± 0.425 |
| **A1S_3026** | | 8.6 ± 0.781 | 3.426 ± .0327 |
| **A1S_3129** | *astB* | 4.066 ± 1.888 | - |

[a] For each strain three independent experiments were performed and represented by the mean ± SD

**Table S5. Mean and standard deviation (SD) of bacterial growth measurement from ATCC 17978 wildtype/mutants and 29D2 wildtype/mutants**

| Locus tag | Gene name | Mean ± SD [OD $_{600\ nm}$][a] | |
|---|---|---|---|
| | | ATCC 17978 | 29D2 |
| **Wildtype** | | 5.87 ± 0.37 | 5.31 ± 0.36 |
| **A1S_2187** | *purH* | 2.16 ± 0.07 | 3.56 ± 0.36 |
| **A1S_2251** | *purF* | 3.38 ± 0.14 | 4.69 ± 0.27 |
| **A1S_2605** | *purM* | 3.35 ± 0.07 | 4.00 ± 0.10 |
| **A1S_2964** | *purE* | 2.45 ± 0.12 | 3.22 ± 0.37 |
| **A1S_1566** | | 4.24 ± 0.08 | 4.41 ± 0.13 |
| **A1S_2687** | *carB* | 0.92 ± 0.008 | 1.03 ± 0.04 |
| **A1S_0414** | | 4.99 ± 0.07 | 5.99 ± 0.07 |
| **A1S_1624** | | 4.72 ± 0.11 | - |
| **A1S_0447** | *rpmG* | 4.56 ± 0.23 | - |
| **A1S_0778** | *metG* | 4.77 ± 0.31 | - |
| **A1S_0530** | | 5.06 ± 0.24 | - |
| **A1S_3366** | *gshA* | 4.68 ± 0.15 | 4.28 ± 0.02 |
| **A1S_1970** | | 4.70 ± 0.27 | 5.96 ± 0.24 |
| **A1S_2840** | *ompA* | 5.19 ± 0.21 | 1.36 ± 0.73 |
| **A1S_3297** | | 4.71 ± 0.20 | - |

| | | | |
|---|---|---|---|
| **A1S_2453** | *ddc* | 3.60 ± 0.14 | 1.90 ± 0.13 |
| **A1S_2454** | *dat* | 3.76 ± 0.14 | 2.99 ± 0.35 |
| **A1S_0113** | | 4.89 ± 0.11 | - |
| **A1S_0116** | | 5.66 ± 0.07 | 5.21 ± 0.10 |
| **A1S_0222** | *aamA* | 4.97 ± 0.06 | 5.62 ± 0.08 |
| **A1S_0065** | *galE* | 4.77 ± 0.12 | 2.48 ± 0.07 |
| **A1S_0806** | | 4.71 ± 0.50 | 5.31 ± 0.94 |
| **A1S_1055** | | 5.91 ± 0.16 | 5.18 ± 0.32 |
| **A1S_2182** | *gidA* | 4.31 ± 0.10 | 5.03 ± 0.24 |
| **A1S_2334** | *sahH* | 4.76 ± 0.05 | - |
| **A1S_2587** | *ruvA* | 4.56 ± 0.04 | - |
| **A1S_2610** | *comEC* | 5.78 ± 0.04 | 3.84 ± 0.09 |
| **A1S_2761** | *prpF* | 4.06 ± 0.11 | 2.12 ± 0.16 |
| **A1S_3026** | | 5.36 ± 0.08 | 5.91 ± 0.25 |
| **A1S_3129** | *astB* | 4.31 ± 0.04 | - |

[a] Bacterial cultures were incubated for 9 hours at 37°C under shaking. For each strain, data obtained from three independent cultures grown on the same day were averaged. In this table endpoint measurements after 9 hours of growth are represented by the mean ± SD

**Table S6. List of p-values for every monitored time point after *Galleria mellonella* caterpillar infection comparing ATCC 17978 wildtype and mutant strains**

| Locus tag | Gene name | p-values[a] | | | | |
|---|---|---|---|---|---|---|
| | | 24 hours p.i. | 48 hours p.i. | 72 hours p.i. | 96 hours p.i. | 120 hours p.i. |
| **A1S_2187** | *purH* | 0.8335 | 0.3506 | 0.3227 | >0.9999 | 0.7110 |
| **A1S_2251** | *purF* | 0.2125 | 0.3506 | 0.3913 | 0.4700 | 0.5708 |
| **A1S_2605** | *purM* | 0.0016 | 0.0050 | 0.0006 | 0.0020 | 0.0166 |
| **A1S_2964** | *purE* | 0.0185 | 0.0524 | 0.0138 | 0.0106 | 0.0698 |
| **A1S_1566** | | 0.0041 | 0.0045 | 0.0006 | 0.0015 | 0.0030 |
| **A1S_2687** | *carB* | <0.0001 | <0.0001 | <0.0001 | <0.0001 | <0.0001 |
| **A1S_0414** | | 0.0005 | 0.0037 | 0.0022 | 0.0048 | 0.0412 |
| **A1S_1624** | | 0.0331 | 0.0306 | 0.0383 | 0.0398 | 0.0835 |
| **A1S_0447** | *rpmG* | 0.0236 | 0.0034 | 0.0334 | 0.0295 | 0.0829 |
| **A1S_0778** | *metG* | <0.0001 | <0.0001 | <0.0001 | <0.0001 | <0.0001 |
| **A1S_0530** | | 0.2037 | 0.0273 | 0.0383 | 0.1546 | 0.1478 |
| **A1S_3366** | *gshA* | 0.3295 | 0.0306 | 0.0049 | 0.0041 | 0.0104 |
| **A1S_1970** | | <0.0001 | 0.0009 | 0.0002 | 0.0003 | 0.0022 |
| **A1S_2840** | *ompA* | <0.0001 | <0.0001 | <0.0001 | <0.0001 | <0.0001 |

| | | | | | | |
|---|---|---|---|---|---|---|
| **A1S_3297** | | 0.0041 | 0.0013 | 0.0022 | 0.0034 | 0.0059 |
| **A1S_2453** | *ddc* | 0.0031 | 0.0590 | 0.0087 | 0.0263 | 0.0829 |
| **A1S_2454** | *dat* | 0.0567 | 0.0537 | 0.0179 | 0.0263 | 0.0306 |
| **A1S_0113** | | 0.0698 | 0.0081 | 0.0021 | 0.0125 | 0.2000 |
| **A1S_0116** | | 0.7542 | 0.0950 | 0.0282 | 0.1064 | 0.1643 |
| **A1S_0222** | *aamA* | <0.0001 | <0.0001 | <0.0001 | 0.0006 | 0.0008 |
| **A1S_0065** | *galE* | <0.0001 | <0.0001 | <0.0001 | <0.0001 | <0.0001 |
| **A1S_0806** | | 0.0185 | 0.2196 | 0.1287 | 0.0876 | 0.1270 |
| **A1S_1055** | | 0.0005 | 0.0002 | <0.0001 | 0.0001 | 0.0012 |
| **A1S_2182** | *gidA* | 0.0016 | 0.1395 | 0.0474 | 0.1778 | 0.1826 |
| **A1S_2334** | *sahH* | 0.1705 | 0.0306 | 0.0053 | 0.0087 | 0.0474 |
| **A1S_2587** | *ruvA* | 0.0063 | 0.0045 | 0.0433 | 0.1036 | 0.2000 |
| **A1S_2610** | *comEC* | <0.0001 | 0.0012 | 0.0005 | 0.0005 | 0.0015 |
| **A1S_2761** | *prpF* | 0.0676 | 0.0335 | 0.0190 | 0.0398 | 0.1643 |
| **A1S_3026** | | 0.7363 | 0.7220 | 0.3227 | >0.9999 | 0.3778 |
| **A1S_3129** | *astB* | 0.0264 | 0.0537 | 0.0422 | 0.0270 | 0.1190 |

[a] Compared to ATCC 17978 WT; unpaired t-test was performed after 24, 48, 72, 96, 120 hours p.i.; p-value ≤ 0.05, *;  p-value ≤ 0.01, **;  p-value ≤ 0.001, ***; p-value ≤ 0.0001, ****

**Table S7. List of p-values for every monitored time point after *Galleria mellonella* caterpillar infection comparing 29D2 wildtype and mutant strains**

| Locus tag | Gene name | p-values[a] | | | | |
|---|---|---|---|---|---|---|
| | | **24 hours p.i.** | **48 hours p.i.** | **72 hours p.i.** | **96 hours p.i.** | **120 hours p.i.** |
| **A1S_2187** | *purH* | 0.0129 | 0.0469 | 0.0657 | 0.0657 | 0.0479 |
| **A1S_2251** | *purF* | >0.9999 | 0.6530 | 0.5072 | 0.5072 | 0.3739 |
| **A1S_2605** | *purM* | 0.0196 | 0.0723 | 0.4512 | 0.5391 | 0.2051 |
| **A1S_2964** | *purE* | 0,0390 | 0,0197 | 0,0403 | 0,0719 | 0.0676 |
| **A1S_1566** | | 0,0046 | 0,0003 | 0,0005 | 0,0011 | 0,0010 |
| **A1S_2687** | *carB* | 0,0012 | <0.0001 | <0.0001 | <0.0001 | <0.0001 |
| **A1S_0414** | | 0.5087 | 0.7318 | 0.3398 | 0.2625 | 0.2275 |
| **A1S_3366** | *gshA* | 0.1630 | 0.1730 | 0.1856 | 0.1898 | 0.1705 |
| **A1S_1970** | | 0.7442 | 0.8998 | 0.8878 | 0.8667 | 0.4248 |
| **A1S_2840** | *ompA* | <0,0001 | <0.0001 | <0.0001 | <0.0001 | <0.0001 |
| **A1S_2453** | *ddc* | 0,0011 | 0,0307 | 0,0274 | 0,0240 | 0,0527 |
| **A1S_2454** | *dat* | 0,0010 | 0,0004 | 0,0152 | 0,0194 | 0,1448 |
| **A1S_0116** | | 0,5020 | 0,1795 | 0,2192 | 0,1486 | 0,1877 |
| **A1S_0222** | *aamA* | 0,1938 | 0,0190 | 0,0176 | 0,0007 | 0,0213 |
| **A1S_0065** | *galE* | 0,0002 | <0.0001 | <0.0001 | <0.0001 | <0.0001 |

| | | | | | | |
|---|---|---|---|---|---|---|
| **A1S_0806** | | 0,0125 | 0,0199 | 0,0092 | 0,0024 | 0,0066 |
| **A1S_1055** | | <0,0001 | <0,0001 | 0,0001 | 0,0003 | 0,0009 |
| **A1S_2182** | *gidA* | 0,0001 | 0,0027 | 0,0155 | 0,0219 | 0,0054 |
| **A1S_2610** | *comEC* | <0,0001 | <0.0001 | <0.0001 | <0.0001 | <0.0001 |
| **A1S_2761** | *prpF* | <0,0001 | <0,0001 | <0,0001 | <0,0001 | <0,0001 |
| **A1S_3026** | | 0,0005 | 0,0003 | 0,0005 | <0,0001 | <0,0001 |

[a] Compared to 29D2 WT; unpaired t-test was performed after 24, 48, 72, 96, 120 hours p.i.; p-value ≤ 0.05, *; p-value ≤ 0.01, **; p-value ≤ 0.001, ***; p-value ≤ 0.0001, ****
